# The relevance of dominance and functional annotations to predict agronomic traits in hybrid maize

**DOI:** 10.1101/745208

**Authors:** Guillaume P. Ramstein, Sara J. Larsson, Jason P. Cook, Jode W. Edwards, Elhan S. Ersoz, Sherry Flint-Garcia, Candice A. Gardner, James B. Holland, Aaron J. Lorenz, Michael D. McMullen, Mark J. Millard, Torbert R. Rocheford, Mitchell R. Tuinstra, Peter J. Bradbury, Edward S. Buckler, M. Cinta Romay

## Abstract

Heterosis has been key to the development of maize breeding but describing its genetic basis has been challenging. Previous studies of heterosis have shown the contribution of within-locus complementation effects (dominance) and their differential importance across genomic regions. However, they have generally considered panels of limited genetic diversity and have shown little benefit to including dominance effects for predicting genotypic value in breeding populations. This study examined within-locus complementation and enrichment of genetic effects by functional classes in maize. We based our analyses on a diverse panel of inbred lines crossed with two testers representative of the major heterotic groups in the United States (1,106 hybrids), as well as a collection of 24 biparental populations crossed with a single tester (1,640 hybrids). We assayed three agronomic traits: days to silking (DTS), plant height (PH) and grain yield (GY). Our results point to the presence of dominance for all traits, but also among-locus complementation (epistasis) for DTS and genotype-by-environment interactions for GY. Consistently, dominance improved genomic prediction for PH only. In addition, we assessed enrichment of genetic effects in classes defined by genic regions (gene annotation), structural features (recombination rate and chromatin openness), and evolutionary features (minor allele frequency and evolutionary constraint). We found support for enrichment in genic regions and subsequent improvement of genomic prediction for all traits. Our results point to mechanisms by which heterosis arises through local complementation in proximal gene regions and suggest the relevance of dominance and gene annotations for genomic prediction in maize.

## INTRODUCTION

Since the development of the first maize hybrids by Shull (1908) and their widespread adoption starting in the 1930s, heterosis has been central to the improvement of maize in the United States. Heterosis, or hybrid vigor, refers to the increase in performance of hybrids relatively to their average parental performance (Shull 1914). There has been little doubt about the practical significance of hybrid vigor as it drove considerable breeding gains in maize during the 20^th^ century, but there has been a long-lasting scientific debate about the basis for this phenomenon (Crow 1998). Predominant hypotheses about the causes of heterosis have related to genetic complementation of parental genomes. The basis for such complementation consists of nonadditive genetic effects, particularly (over)dominance (within-locus complementation, i.e., interaction between alleles within single genetic loci) and epistasis (among-locus complementation, i.e., interactions involving multiple genetic loci). Overdominance, or heterozygous advantage, was initially favored as an explanation for heterosis (East 1936, Crow 1948). However, this type of gene action did not account for experimental results, such as the decrease in the realized degree of dominance over consecutive generations in populations derived from biparental crosses (Gardner 1963, Moll et al. 1964). Instead, it was proposed that apparent overdominance was due to dominance gene action at closely-linked polymorphisms having opposite effects (repulsion phase linkage) (Hill and Robertson 1966, Cockerham and Zeng 1996, Graham et al. 1997). Epistasis also provides a plausible explanation for genomic complementation. However, studies assessing its contribution to heterosis have suffered from a lack of statistical power (Reif et al. 2005) and have reported contrasting results (e.g., Mihaljevic et al. 2005 and Ma et al. 2007).

Genetic studies in maize have investigated dominance gene action by focusing either on directional dominance, effects of quantitative trait loci (QTL), or genome-wide (polygenic) effects. Studies on testcrosses or diallel mating designs have investigated directional dominance by assessing the relationship between heterosis and inter-parent genetic distance (e.g., Reif et al. 2003), or the relationship between testcross means and the genomic contribution of a given parent to the testcross (e.g., Hinze and Lamkey 2003). Their conclusions seem to support the presence of directional dominance, particularly for grain yield. Furthermore, studies on populations derived from backcrosses between recombinant inbred lines and their parents, under North Carolina III designs, have generally identified several QTL with significant dominant effects for traits such as flowering time, plant height, and grain yield (e.g., Frascaroli et al. 2007, Larièpe et al. 2012). Finally, genomic prediction analyses in maize have assessed polygenic dominance effects for their contribution to genotypic variability. Importantly, these genomic prediction studies have often focused on factorial designs in which hybrids were obtained from crosses between lines coming from different heterotic groups: Flint and Dent (e.g., Technow et al. 2014) or Stiff Stalk and non-Stiff Stalk (e.g., Kadam et al. 2016). Most of these studies have suggested little contribution of non-additive effects (i.e., specific combining abilities) to genotypic variability. However, they could not assess the relevance of dominance effects in more diverse panels in which genomic effects, and heterotic responses, may be more inconsistent, due to differential levels of genomic complementation within and across heterotic groups (Reif et al. 2005, Gerke et al. 2015).

The above-mentioned studies have assayed the relative importance of additive and dominance effects across the genome, but they have not attempted to describe the properties of genomic regions most enriched for causal variants. Other studies in maize have characterized the genetic basis of agronomic traits based on locus properties such as gene proximity, structural features, and/or evolutionary features. Gene proximity has been linked to causal variants in maize through enrichment for QTL effects (Wallace et al. 2014); additionally, a large portion of variability of gene expression in maize has been attributed to *cis* polymorphisms (Schadt et al. 2003). Therefore, most polymorphisms underlying genome complementation and hybrid vigor are expected to lie in proximal gene regions. Structural features may also be functionally relevant to heterosis in maize. For example, chromatin openness and high recombination rate were associated with enrichment for QTL effects in maize inbred lines (Rodgers-Melnick et al. 2016). However, studies on maize hybrids have also shown that heterotic QTL tend to locate around centromeres, where recombination rate is low (Larièpe et al. 2012, Thiemann et al. 2014, Martinez et al. 2016). Therefore, it is possible that causal loci for hybrid vigor in maize is enriched in regions characterized by low recombination rate and closed chromatin, because of repulsion phase linkage (Hill and Robertson 1966). Evolutionary features characterize allelic diversity within species (e.g., allele frequency or nucleotide diversity) and across species (e.g., evolutionary constraint). Lower allelic diversity has been associated with stronger QTL effects in hybrid maize (Mezmouk and Ross-Ibarra 2014, Yang et al. 2017). Therefore, loci with low allele frequency or high evolutionary constraint may have stronger effects on heterosis in maize. Importantly, structural and evolutionary features have also been associated with gene density. For example, Beissinger et al. (2016) and Rodgers-Melnick et al. (2016) have reported lower nucleotide diversity and more open chromatin near genes, respectively. So, there is ambiguity about the relevance of evolutionary and structural features to capture variability at agronomic traits independently from gene proximity.

In this study, we aimed at characterizing the genetic basis of hybrid vigor for three agronomic traits (days to silking, plant height, and grain yield) in panels representative of genetic diversity in maize. We analyzed two hybrid panels: one was derived from crosses between a diverse sample of maize inbred lines and either of two testers, B47 and PHZ51, belonging respectively to the Stiff Stalk (SS) and non-Stiff Stalk (NSS) heterotic groups; the other was derived from crosses between the US Nested Association Mapping (NAM) panel and PHZ51. We investigated the importance of dominance for heterosis in maize by (i) the contribution of polygenic dominance to genotypic variability, (ii) the existence of significant dominance effects at QTL, and (iii) directional effects of dominance by inbreeding. In addition, we tested the hypotheses that most genetic effects involved in dominance are located (i) near genes, (ii) in low-recombination regions, and (iii) at evolutionarily constrained loci (Figure 1). Our study is focused on the usefulness of genetic effects partitioned by gene action (additive or dominance effects) and functional classes (based on gene proximity and structural or evolutionary features), for applications such as prioritization of SNP markers and genomic prediction.

**Figure 1.**
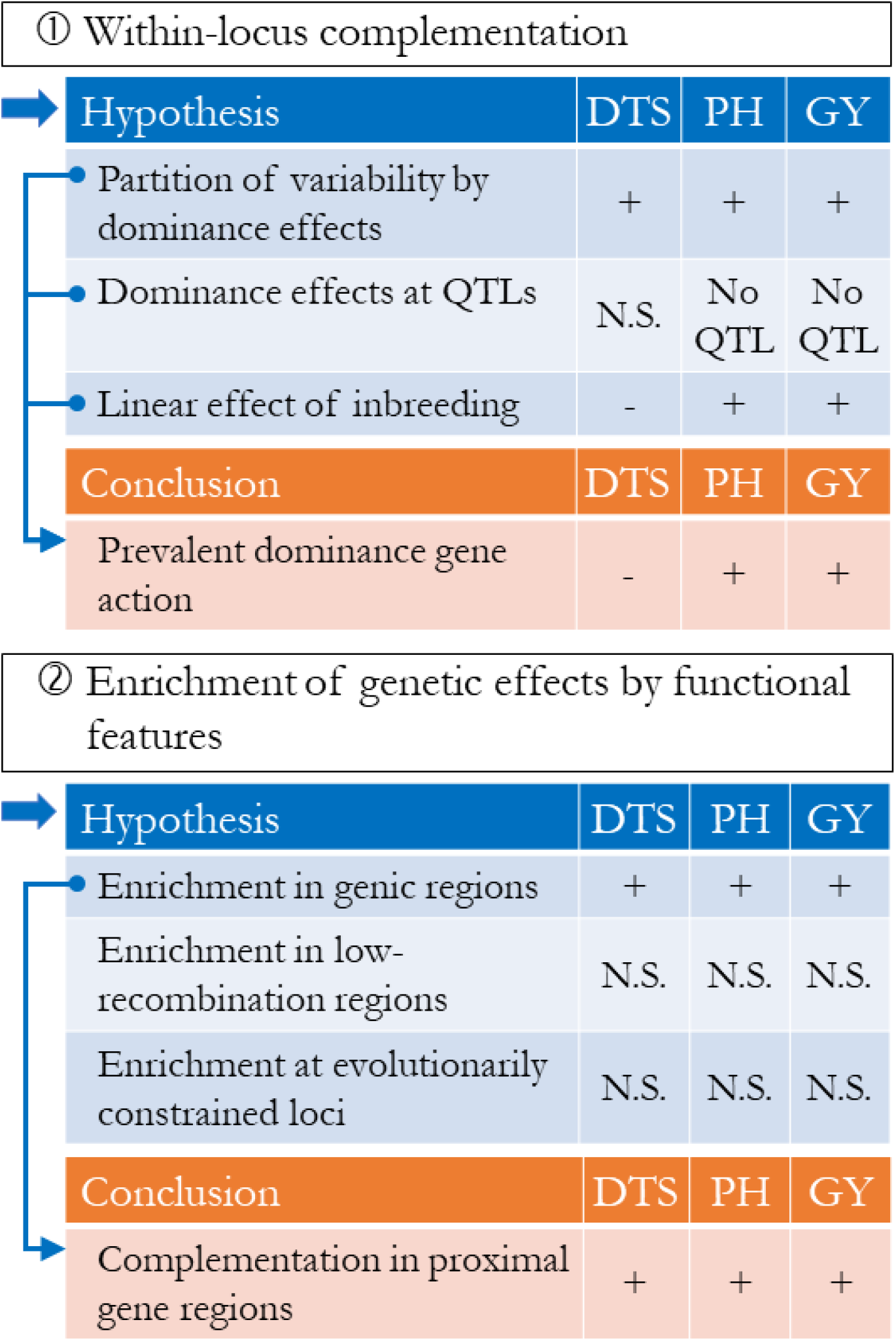
Graphical summary of the study. Two rationales were tested: (1) dominance gene action explains heterosis in maize; (2) genetic effects underlying heterosis are enriched by functional classes. Under each rationale, evidence from analyses is characterized as consistent (+) or inconsistent (-) with scientific hypotheses. Non-conclusive evidence is either due to absence of QTL (No QTL) or lack of significance (N.S.).

## MATERIAL AND METHODS

### Phenotypic data

#### Phenotypic measurements

In this study, two panels of maize lines were assayed for hybrid performance: the NCRPIS association panel (hereafter, Ames) and the nested association mapping panel (hereafter, NAM). The Ames panel comprises a subset of temperate inbred lines from the diversity panel described by Romay et al. (2013); the NAM panel is a subset of 24 recombinant inbred line (RIL) populations, all having one parent in common, B73, as described by McMullen *et al.* (2009).

In the hybrid Ames panel, a subset of 875 inbred lines was selected to reduce differences in flowering time while favoring genetic diversity based on pedigree information. Two inbred lines, formerly under Plant Variety Protection, were selected as testers: one non-Stiff Stalk (NSS) inbred (PHZ51) and one Stiff Stalk (SS) inbred (B47, also known as PHB47). Inbreds were assigned to one or two testers based on known heterotic group: SS inbreds were crossed with PHZ51, while NSS inbreds were crossed with B47; inbreds with unknown heterotic group as well as inbreds belonging to the Goodman association panel (Flint-Garcia et al. 2005) were crossed with both testers, for a total of 1,111 hybrids. Hybrids were assigned to one of four combinations, based on tester (PHZ51 or B47) and maturity (early or late). Each combination was split into three sets based on expected plant height (short, medium, or tall). Each of those 12 groups were arranged in an incomplete block (alpha-lattice) design. Sets were randomized for each environment, and tester-maturity combinations were randomized within each set. One common check (B73×PHZ51) was randomly included in each block of the lattices, and each lattice randomly included three additional checks (PHZ51×B47, B47×PHZ51, and a maturity commercial check). In the Ames panel, evaluation was performed in 2011 and 2012, in six locations across the US – Ames (IA), West Lafayette (IN), Kingston (NC), Lincoln (NE), Aurora (NY), and Columbia (MO) – for a total of nine unique environments: 11IA, 11IN, 11NC, 11NE, 11NY, 11MO, 12NE, 12NC, and 12MO.

In the hybrid NAM panel, selection and evaluation were performed as described by Larsson et al. (2017). Briefly, a subset of 60 to 70 RILs from each of the NAM families was selected to reduce differences in flowering time across families: the later RILs from the earliest families and the earlier RILs from the latest families, for a total of 1,799 RILs. All RILs were crossed with the same tester: PHZ51. Hybrids were evaluated in five different locations – Ames (IA), West Lafayette (IN), Kingston (NC), Aurora (NY), and Columbia (MO) – during 2010 and 2011 for a total of eight unique environments: 10IA, 10IN, 10NC, 10MO, 11IA, 11IN, 11NC, and 11NY.

Both NAM and Ames hybrids were planted in two-row plots (40-80 plants per plot; 50,000 to 75,000 plants per hectare), except for 11NY, where 12 plants were planted per plot. The following traits were measured: days to silking (number of days from planting until 50% of the plants had silks; DTS), plant height (cm from soil to flag leaf; PH), and grain yield (t/ha adjusted to 15.5% moisture; GY). In 11NY, only PH and DTS were measured (Table 1).

#### Genotype means and heritability

Genotype means of hybrids were estimated by a linear mixed model, fitted by ASREML-R v3.0 (Butler et al. 2009). For each combination of panel (Ames or NAM panel) and trait (DTS, PH, or GY), the following effects were estimated: genotype [fixed], environment [random, independent, and identically normally distributed (i.i.d.)], field within environment [random, i.i.d.], and, if possible, spatial effects within environment/field combinations [random, normally distributed under first-order autoregressive covariance structures by row and column]. Since genotypes were not replicated within environments, genotype-by-environment interactions were pooled with residual variation. For PH in both panels, spatial effects were not included in the model because the fitting algorithm could not converge to a solution. For GY in both panels, DTS measurements [fixed] were included in the model to account for phenological differences among lines. In addition to estimating genotype effects as fixed, models with genotype effects as random were also fitted to estimate genotypic variance 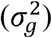 and error variance 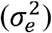. Broad-sense heritability on a plot basis was then calculated as 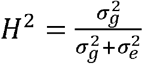. Finally, entry-mean reliability was estimated as 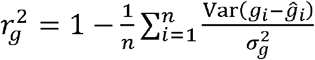 where *n* is the number of hybrids assayed in either panel and 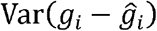 is the prediction error variance of genotype mean for hybrid *i* (Searle et al. 2009).

### Genotypic data

#### Marker data

All the inbreds that were used to create the evaluated hybrids were originally genotyped using genotyping-by-sequencing (GBS) (Romay et al. 2013, Rodgers-Melnick et al. 2015). Singlenucleotide polymorphisms (SNPs) were called with the software TASSEL v5.0 (Bradbury et al. 2007) using the GBS production pipeline and the ZeaGBSv2.7 Production TOPM obtained from more than 60,000 Zea GBS samples (Glaubitz et al. 2014).

The GBS SNPs in both panels were used for imputing marker scores (alternate-allele counts) called at whole-genome-sequencing (WGS) SNPs from the Hapmap 3.2.1 panel, under version 4 of the reference B73 genome (Bukowski et al. 2018). From the original WGS dataset heterozygote SNPs were set to missing (since these were presumably due to errors or collapsed paralogous loci) and WGS SNPs were filtered out if they did not satisfied the following criteria: two alleles by SNP, call rate > 50%, and minor allele count > 3. A total of 25,555,019 positions across the reference genome were then selected for imputation. Marker scores were imputed by BEAGLE v5 (Browning and Browning 2018), with the following parameters: 10 burn-in iterations, 15 sampling iterations, and effective population size set to 1000. Marker scores at WGS SNPs were first fully imputed and phased in the Hapmap 3.2.1 panel; then, they were imputed in the Ames panel and the NAM panel separately, based on GBS SNPs using the imputed Hapmap 3.2.1 panel as reference.

In subsequent analyses, hybrids were divided in four sets: Ames/PHZ51, Ames/B47, the entire Ames hybrid panel (Ames/PHZ51+B47), and NAM/PHZ51. These sets comprised 463, 643, 1106, and 1640 hybrids, respectively. After imputation, WGS SNPs were further filtered for the following criteria in every set, based on the respective subsets of inbreds: minor allele frequency ≥ 0.01; estimated squared correlation between imputed and actual marker scores ≥ 0.8 (Browning and Browning 2009). Marker scores at selected WGS SNPs were then inferred for each hybrid by using CreateHybridGenotypesPlugin in TASSEL v5.0; at each selected WGS SNP, female and tester marker scores were combined, unless either of these was heterozygous or missing (in which case the hybrid genotype was set to missing). After filtering by quality and variability of marker scores, a total of *m* = 12,659,487 WGS SNPs were retained for subsequent analyses (14,846,984 to 15,733,697 SNPs were selected due to filters on minor allele frequency alone). In a given set, the marker data consisted of the matrix **X** of minor-allele counts, where minor alleles were defined by frequencies in the Hapmap 3.2.1 panel, and the matrix **Z** of heterozygosity, which coded homozygotes as 0 and heterozygotes as 1.

#### Population principal components

Principal component analysis (PCA) was performed using the R package irlba v2.3.3 (Baglama and Reichel 2005), based on the Goodman association panel, presumed to represent the genetic diversity among elite maize inbred lines (Flint-Garcia et al. 2005). Matrix **P**, consisting of coordinates at the first three PCs in hybrids, was obtained by (i) adjusting marker scores by their observed mean in the Goodman association panel, and (ii) mapping adjusted marker scores to PCs by the SNP loadings from PCA, i.e., **P** = (**X** – **M**)**V**, where **X** – **M** is the matrix of adjusted marker scores and **V** is the *m*×3 matrix of right-singular vectors from PCA.

### Functional features

#### Gene annotation: proximity to genes

Gene positions were available from v4 gene annotations, release 40 (ftp://ftp.ensemblgenomes.org/pub/plants/release-40/gff3/zea_mays/Zea_mays.AGPv4.40.gff3.gz). Gene proximity bins (either ‘Proximal’ or ‘Distal’) then indicated whether any given SNP was within 1 kb of an annotated gene (less than 1 kb away from the start or end positions).

#### Structural features: recombination rate and chromatin openness

Previously published recombination maps identified genomic segments originating from either parent within the progeny of each NAM family (Rodgers-Melnick et al. 2015). These maps were uplifted to version 4 of the reference genome using CrossMap v0.2.5 (Zhao et al. 2014). Then, the average numbers of recombination events (recombination fractions) were fitted on genomic positions by a thin-plate regression spline model, by the R package mgcv v1.8-27 (Wood 2003). Based on this model, recombination rates **c** were inferred by finite differentiation of fitted recombination fractions: 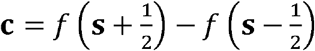, where **s** is the vector of genomic positions of all WGS SNPs, and *f* is the function inferred by the spline model. Finally, we defined recombination bins as follows: *c_j_* ≤ 0.45 cM/Mb, 0.45 cM/Mb < *c_j_* ≤ 1.65 cM/Mb, and 1.65 cM/Mb < *c_j_*, where 0.45 cM/Mb and 1.65 cM/Mb are the first two tertiles of estimated recombination rates *c_j_* among all WGS SNPs.

Chromatin accessibility was previously assessed by micrococcal nuclease hypersensitivity (MNase HS) in juvenile root and shoot tissues in B73 (Rodgers-Melnick et al. 2016). Here, MNase HS peaks were mapped to their coordinates in version 4 of the reference genome. A given SNP was considered to lie in a euchromatic (open) region if a MNase HS peak was detected, in either root or shoot tissues. We then defined MNase HS bins as ‘Dense’ or ‘Open’ for the absence or presence of MNase HS peaks, respectively.

#### Evolutionary features: minor allele frequency and evolutionary constraint

Minor allele frequencies (MAF) at SNPs were determined based on the Hapmap 3.2.1 panel in version 4 of the reference genome, without imputation of marker scores. Similarly to Evans et al. (2018), we defined MAF bins as follows: MAF ≤ 0.01, 0.01 < MAF ≤ 0.05, and 0.05 < MAF (SNPs were not binned at MAF ≤ 0.0025 due to only 7,202 of them falling into this class).

Evolutionary constraints at SNPs were reflected by genomic evolutionary rate profiling (GERP) scores, as introduced by Davydov et al. (2010). Here we derived GERP scores from a whole-genome alignment of 13 plant species (Rodgers-Melnick et al. 2015, Yang et al. 2017), based on coordinates in version 4 of the reference genome. We defined GERP score bins as GERP ≤ 0 and GERP > 0.

### Genome-wide polygenic models

#### Additive effects

Genome-wide additive effects were estimated under a standard genomic BLUP (GBLUP) model (VanRaden 2008), as follows:

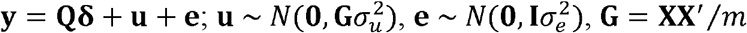

where **y** was the vector of genotype means; **Q** = [**1 P**] was the matrix consisting of a vector of ones and the first three PCs as described above; **δ** were fixed effects; **u** and **e** consisted of polygenic additive genomic effects and random errors, respectively. The GBLUP model was fitted in Ames/PHZ51+B47 or NAM/PHZ51, by restricted maximum likelihood (REML) using the R package regress v1.3-15 (Clifford and McCullagh 2005).

For comparison to Bayesian sparse linear mixed models (see next section below), we also fitted RR-BLUP models where the effects of PCs were not explicitly accounted for by fixed effects, i.e., 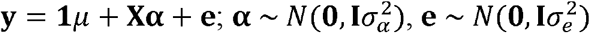 where **α** consisted of random additive marker effects. The RR-BLUP model was fitted in Ames/PHZ51+B47 or NAM/PHZ51, by REML using GEMMA v0.98.1 (Zhou and Stephens 2012).

#### Additive and dominance effects

To account for dominance, the GBLUP model was extended to the dominance GBLUP (DGBLUP) model, as follows:

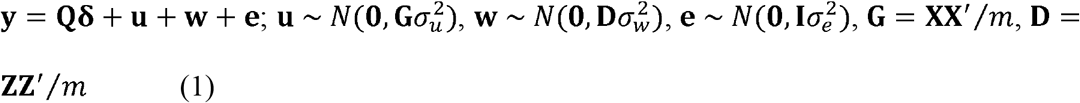

where **w** consisted of polygenic dominance effects. Model (1) was fitted in Ames/PHZ51+B47, by REML using the R package regress v1.3-15 (Clifford and McCullagh 2005).

#### Directional effects

Directional effects arise from consistent genetic effects across loci, such that their average is non-zero. An example of directional effects about dominance is inbreeding depression, due to genome-wide dominance effects being usually positive for fitness. Under a simple dominance model without linkage nor epistasis, inbreeding depression is characterized by a linear negative relationship between the inbreeding coefficient and fitness (Falconer and Mackay 1996). Moreover, in presence of directional epistatic effects, the relationship between the inbreeding coefficient and fitness is expected to be nonlinear (Crow and Kimura 1970). To capture such nonlinearity, specifically dominance×dominance epistasis, the quadratic effect of the inbreeding coefficient was fitted along with its linear effect. We followed Endelman and Jannink (2012) to estimate genomic inbreeding coefficients with respect to a base population, here represented by the Goodman association panel. For each hybrid *i*, the coefficient of genomic inbreeding was calculated as 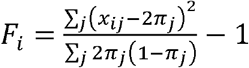, where *π_j_* was the allele frequency in the Goodman association panel.

Directional effects of inbreeding were assayed as fixed effects under an extension of the DGBLUP model (1). The following model was fitted:

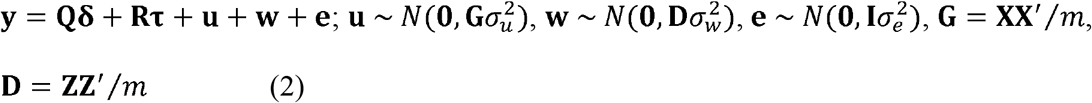

where **R** and **τ** consisted of genomic inbreeding values and their directional effects (linear or quadratic), respectively. Significance of estimates of **τ** was assessed by Wald tests. Model (2) was fitted in Ames/PHZ51+B47 or NAM/PHZ51, by REML using the R package regress v1.3-15 (Clifford and McCullagh 2005).

### Oligogenic models

Oligogenic effects of SNPs were inferred using association models which estimated the effect of each SNP while accounting for background polygenic SNP effects. Two types of models were used: standard linear mixed models, where the effect of each SNP was estimated separately, and Bayesian linear models, where effects of all SNPs under assay were fitted simultaneously.

#### Genome-wide association models

Standard linear mixed models were genome-wide association study (GWAS) models fitted to assess the significance of SNPs for additive effects only (marginal additive effects), or additive and dominance effects simultaneously.

For assessing marginal additive effects (*β_j_*, fixed, for each SNP *j*), the following model was fitted in Ames/PHZ51+B47 or NAM/PHZ51: 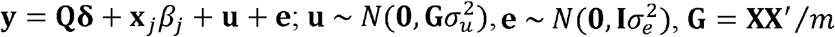. For assessing additive and dominance effects (*β_j_* and *θ_j_*, fixed, for each SNP *j*), the previous model was extended in Ames/PHZ51+B47 to incorporate dominance for both fixed effects and random effects: 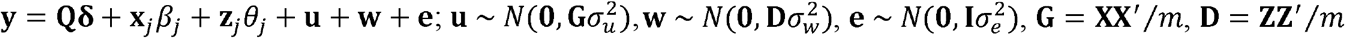. GWAS models were fitted under the EMMAX approximation of Kang et al. (2010), using function fastLm in the R package RcppEigen v0.3.3.5.0 (Bates and Eddelbuettel 2013). Significance of SNPs was assessed by Wald tests on estimates of *β_j_* and *θ_j_*. False discovery rates (FDR) were estimated based on p-values from Wald tests by the method of Benjamini and Hochberg (1995).

#### Bayesian sparse linear mixed models

Models used for joint estimation of additive marker effects were Bayesian sparse linear mixed models (BSLMM) where marker effects are decomposed into a polygenic component and a sparse component (characterizing outstanding effects of few markers). Using Markov chain Monte Carlo (MCMC), the following model was fitted:

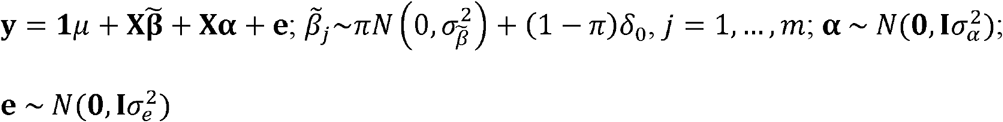

BSLMMs were fitted in Ames/PHZ51+B47 or NAM/PHZ51 by GEMMA v0.98.1, with 1,000,000 and 10,000,000 MCMC iterations for burn-in and sampling, respectively (Zhou et al. 2013). As part of the MCMC process, a vector **γ** of posterior inclusion probabilities (PIP) was generated, such that 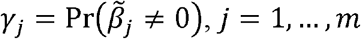. We estimated window posterior inclusion probability (WPIP) following Guan and Stephens (2011), by summing *γ_j_*’s in 500-kb windows, sliding by 250-kb steps.

### Functional polygenic models

#### Effects of markers by evolutionary and structural features

Effects of evolutionary and structural features on the amplitude of marker effects were captured by linear mixed models which partitioned the genomic variance among hybrids by annotation bin. For each feature (gene proximity, recombination rate, chromatin openness, MAF, and GERP) the following model was fitted:

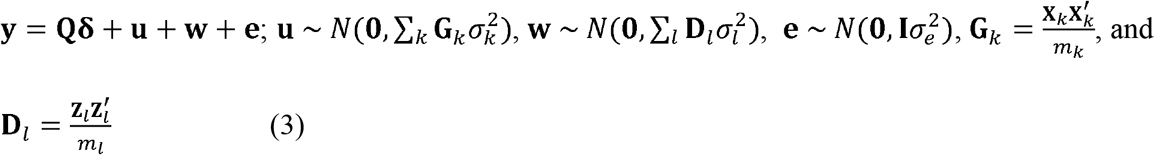

where **X**_*k*_ (**Z**_*l*_) is the matrix of minor-allele counts at the *m_k_* (*m_l_*) SNPs in bin *k* (*l*), and 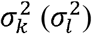 is the variance component associated to additive effects in bin *k* (dominance effects in bin *l*). The significance of the variance partition was assessed by a likelihood ratio test, comparing the REML of the evaluated model to that of a baseline model. Two types of variance partition were analyzed by model (3): partition by one feature (baseline: DGBLUP in Ames/PHZ51+B47 and GBLUP in NAM/PHZ51), and partition by both gene proximity and another feature (baseline: partition by gene proximity only). Model (3) was fitted in Ames/PHZ51+B47 or NAM/PHZ51, by REML using the R package regress v1.3-15 (Clifford and McCullagh 2005).

#### Variance partition and SNP enrichment

For each hybrid *i*, the proportion of variance explained by marker effects in GBLUP was estimated by 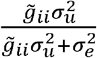, where 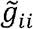 is the *j*^h^ diagonal element of matrix **G** adjusted for fixed effects, i.e., 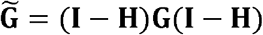, with **H** = **Q**(**Q′Q**)^-1^**Q**′ being the matrix of projection onto the column space of **Q**. The proportion of variance explained by additive marker effects in DGBLUP was estimated by 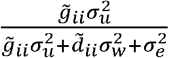, and similarly for dominance effects: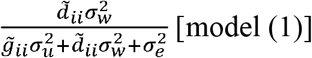. Finally, in functional polygenic models, the proportion of variance explained by additive marker effects at bin *k*\* was estimated by 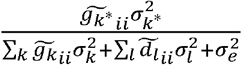, and similarly for dominance effects at bin 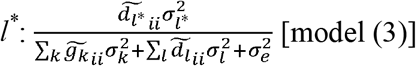. Proportions of variance in whole panels for a given type of effects were then obtained by averaging estimated proportions over hybrids. In functional polygenic models, SNP enrichment for additive effects at bin *k*\* was calculated by the ratio of 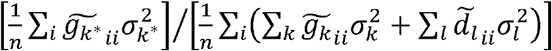, i.e., the proportion of genomic variance explained by bin *k*\*, over *m*_*k*^*^_/[Σ_*k*_ *m_k_* + Σ_*l*_*m_l_*], i.e., the proportion of SNPs in bin *k*\* (and similarly for dominance effects at bin *l*).

### Validation of prediction models in NAM/PHZ51

Models fitted in Ames/PHZ51+B47 were assessed for prediction accuracy (Pearson correlation between observed genotype means and their predicted values) in NAM/PHZ51. Our validation scheme was meant to reflect the merit of prediction models in practical applications of genomic selection, so prediction accuracies were estimated separately in each NAM/PHZ51 population. Therefore, prediction accuracy for any prediction model (e.g., DGBLUP) could be tested for significance of average prediction accuracy (non-zero mean, by a one-sample t-test) and estimated difference in accuracy compared to another model (non-zero difference, by a two-sample t-test paired by population) over NAM/PHZ51 populations.

### Assessment of genotype-by-panel interactions

Interactions between genotypes and panels (environments) were assessed by Pearson correlation in genotypes means between panels, for hybrids which were common to both panels (*ρ_C_*). These hybrids were derived from crosses between PHZ51 and one of 23 check genotypes (B73, B97, CML52, CML69, CML103, CML228, CML247, CML277, CML322, CML333, Il14H, Ki3, Ki11, M162W, M37W, Mo17, Mo18W, NC350, NC358, Oh43, P39, Tx303, and Tzi8).

Genotype-by-panel interactions were also assessed by the following polygenic model, based on Jarquín et al. (2014):

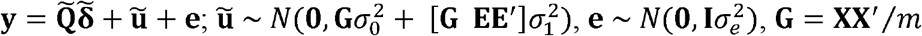

where **E** was the *n*×2 design matrix attributing genotypes to panels (environments), either Ames/PHZ51+B47 or NAM/PHZ51; 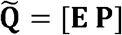 and 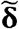 captured effects of panels and population structure; 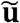 were polygenic genomic effects with main variance and panel-specific variance being quantified by 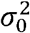 and 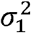, respectively; refers to the Hadamard (element-wise) product. For a given hybrid *i*, correlation in 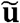 between different panel *j* and *j*’ was defined by 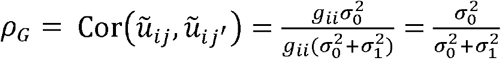 (Jarquín et al. 2014). This model was fitted in Ames/PHZ51+B47 and NAM/PHZ51, by REML using the R package regress v1.3-15 (Clifford and McCullagh 2005).

## RESULTS

### Hybrid panels differed by their genetic diversity and their genetic basis for grain yield

#### Hybrid panels displayed contrasting levels of diversity

The genotypic variability in Ames/PHZ51 and Ames/B47 was well represented by the diversity in the Goodman association panel (Flint-Garcia et al. 2005) (Figure 2). The entire Ames hybrid panel (Ames/PHZ51+B47) involved hybrids with some affinity to semi-tropical lines (e.g., CML 247) but, for the most part, it comprised hybrids closely related to SS lines like B73 and NSS lines like Mo17 (Figure 2). Compared to Ames/PHZ51+B47, NAM/PHZ51 was less diverse, as its genetic composition was relatively consistent (Figure 2). Indeed, NAM/PHZ51 was produced by crosses between a single NSS tester (PHZ51) and bi-parental populations which were all derived from a cross involving B73 as a common parent (i.e., NAM RILs are 50% B73). Moreover, female parents in NAM/PHZ51 were selected for similar flowering time to PHZ51, hence narrowing down further the genetic diversity in this panel.

**Figure 2.**
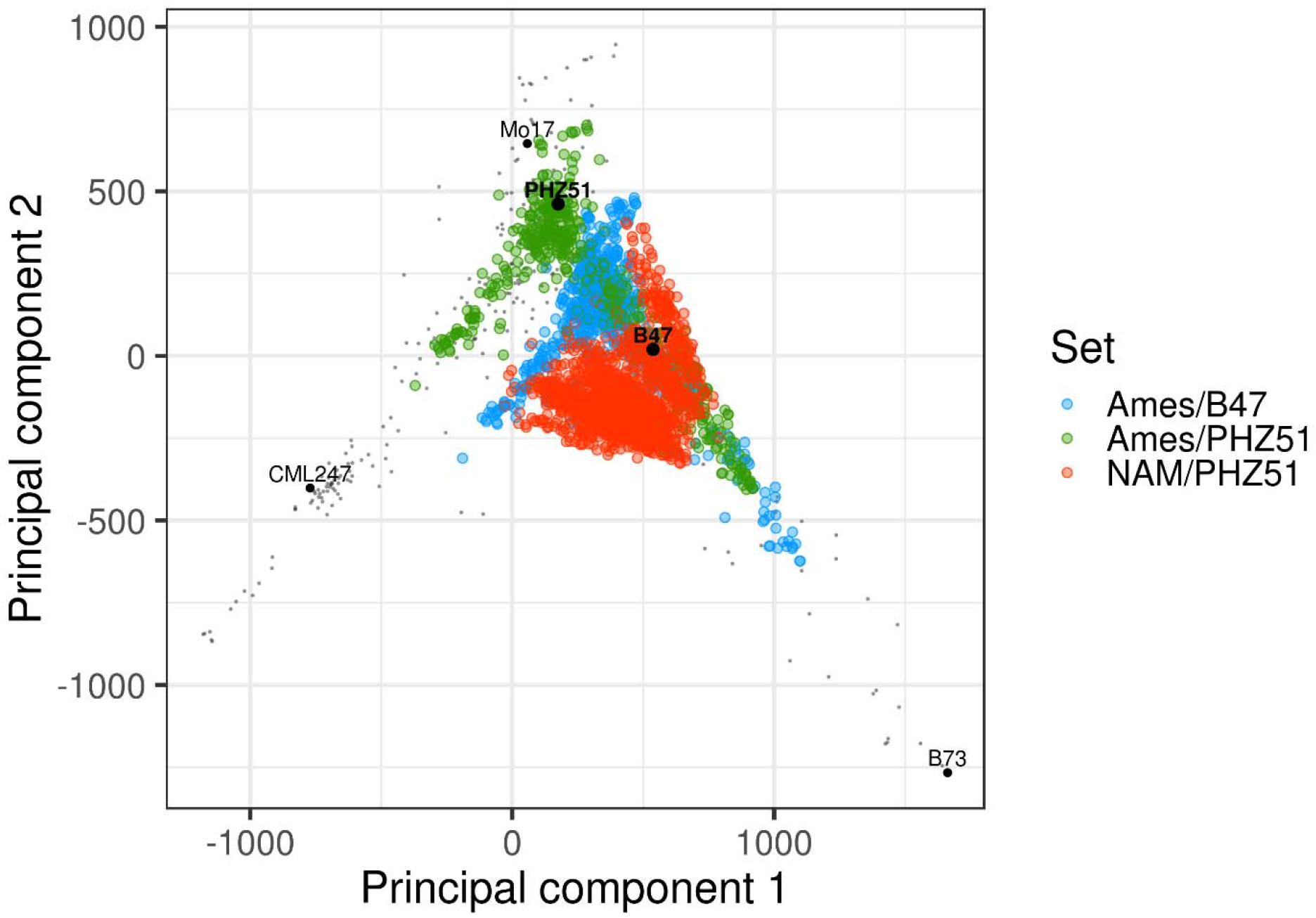
The two hybrid panels differ by their number of testers and their level of diversity. Principal component analysis (PCA) plot of hybrids, by set. Black dots refer to inbred lines in the Goodman association panel (Flint-Garcia et al. 2005), a subset of the Ames panel. B73: SS reference line; Mo17: NSS reference line; CML247: CIMMYT semi-tropical reference line.

#### Genome-wide patterns across panels were similar for linkage disequilibrium but not for allele frequency

Linkage disequilibrium (LD) patterns were quite similar in both hybrid panels. After adjustment for population structure and relatedness (following Mangin et al. 2012), LD values were very concordant between Ames/PHZ51 and Ames/B47 (*r*=0.95), and fairly concordant between Ames/PHZ51+B47 and NAM/PHZ51 (*r*=0.77) (Figure S1). Average LD values along chromosomes decayed at similar rates, reaching 0.1 at 160 kb in Ames/PHZ51+B47 and 151 kb in NAM/PHZ51. However, despite relatively fast LD decay, variance in LD values over SNP pairs was large (Figure S1). Allele frequencies among female parents were very concordant between Ames/PHZ51 and Ames/B47 (*r*=0.98), and fairly concordant between Ames/PHZ51+B47 and NAM/PHZ51 (*r*=0.88) (Figure S2). However, for a subset of markers, frequency spectra were clearly dissimilar, since SNPs at relatively low frequency in NAM/PHZ51 (< 0.5) had frequencies between 0 and 1 in Ames/PHZ51+B47 (Figure S2). Such differences in allele frequency may result in inconsistencies in genetic effects across panels because of dominance and epistatic interactions (Mäki-Tanila and Hill 2014).

#### Genetic bases for grain yield were inconsistent across panels

Three agronomic traits were analyzed for heterosis in Ames/PHZ51+B47 and NAM/PHZ51: days to silking (DTS), plant height (PH), and grain yield adjusted for differences in flowering time among hybrids (GY). The relatively low accuracy of genotype means for GY (as reflected by low broad-sense heritability and entry-mean reliability; Table 1) suggested variability due to genotype-by-environment interactions. Accordingly, genotypic effects appeared highly inconsistent for GY between Ames/PHZ51+B47 and NAM/PHZ51 (Table 2). For GY, correlations across panels based on genotype means of checks (*ρ_C_*) and genomic marker effects (*ρ_G_*) were not significantly different from zero (*p* > 0.10; Table 2). In contrast, consistency in genetic bases was higher for PH (*ρ_C_* = 0.65, *ρ_G_* = 0.78; *p* < 0.001) and DTS (*ρ_C_* = 0.93, *ρ_G_* = 1.0; *p* < 0.001) (Table 2). Although *p_G_* may reflect interactions with genetic backgrounds across panels, *ρ_C_* merely assessed consistency in the performance of identical checks across panels, reflecting only differences between environments (locations, years, management regimens, etc.). Because *ρ_C_* and *ρ_G_* were generally concordant, marker-by-panel interactions, as quantified by both *ρ_C_* and *ρ_G_*, likely reflected sensitivity of marker effects to environments. Therefore, DTS, PH, and GY would represent three distinct levels of sensitivity to genotype-by-environment interactions, being respectively weak, moderate, and strong.

**Table 1.** Phenotypic information by panel and trait

| Panel | Trait | Environments | Mean | H<sup>2</sup> | $r_g^2$ |
| --- | --- | --- | --- | --- | --- |
| Ames | DTS | 11IA 11IN 11NC 11NE 11NY 11MO<br>12NC 12NE 12MO | 66.4 | 0.78 | 0.95 |
|  | PH | 11IA 11IN 11NC 11NE 11NY 11MO<br>12NC 12NE 12MO | 219 | 0.69 | 0.92 |
|  | GY | 11IA 11NC 11NE 11MO<br>12NC 12NE 12MO | -0.02 | 0.29 | 0.62 |
| NAM | DTS | 10IA 10IN 10MO<br>11IA 11IN 11NC 11NY | 70.6 | 0.55 | 0.81 |
|  | PH | 10IA 10IN 10NC 10MO<br>11IA 11IN 11NC 11NY | 247 | 0.30 | 0.69 |
|  | GY | 10IA 10IN 10MO<br>11IA 11NC | -0.20 | 0.16 | 0.35 |

Trait: days to silking (DTS), plant height (PH), grain yield adjusted for DTS (GY). Environments refer to year (2010, 2011, 2012) and locations [Kingston (NC), Ames (IA), West Lafayette (IN), Lincon (NE), Columbia (MO), and Aurora (NY)]. Mean: average phenotypic value. H<sup>2</sup>: broad-sense heritability on a plot basis. $r_g^2$ : average entry-mean reliability.

**Table 2.**
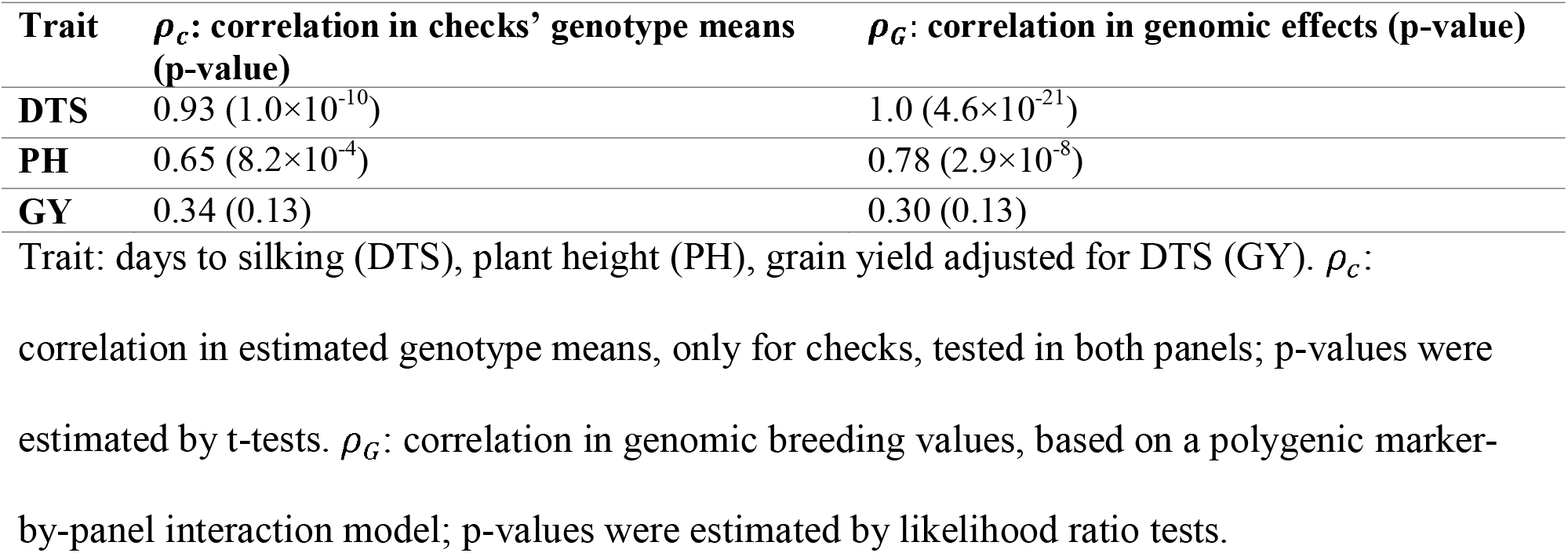
Interactions between genotypes and environments/panels

### Heterosis for plant height and grain yield appeared to be caused by dominance gene action

#### Polygenic dominance effects captured genotypic variability for all traits

To assess the general relevance of polygenic dominance effects, genotypic variability captured in our assay was partitioned into additive and dominance components in a dominance GBLUP (DGBLUP) model. For all traits, dominance accounted for a significant portion of genotypic variability in Ames/PHZ51+B47 (*p* ≥ 2.2×10^-11^), capturing 35%, 23%, and 41% of genomic variance for DTS, PH, and GY (Figure 3a). These estimates corresponded to average degrees of dominance (ratio of dominance-to-additive standard deviations) of 0.73, 0.54, and 0.83 respectively. Therefore, overdominance did not seem to be pervasive in Ames/PHZ51+B47 (average degrees of dominance lower than one).

**Figure 3.**
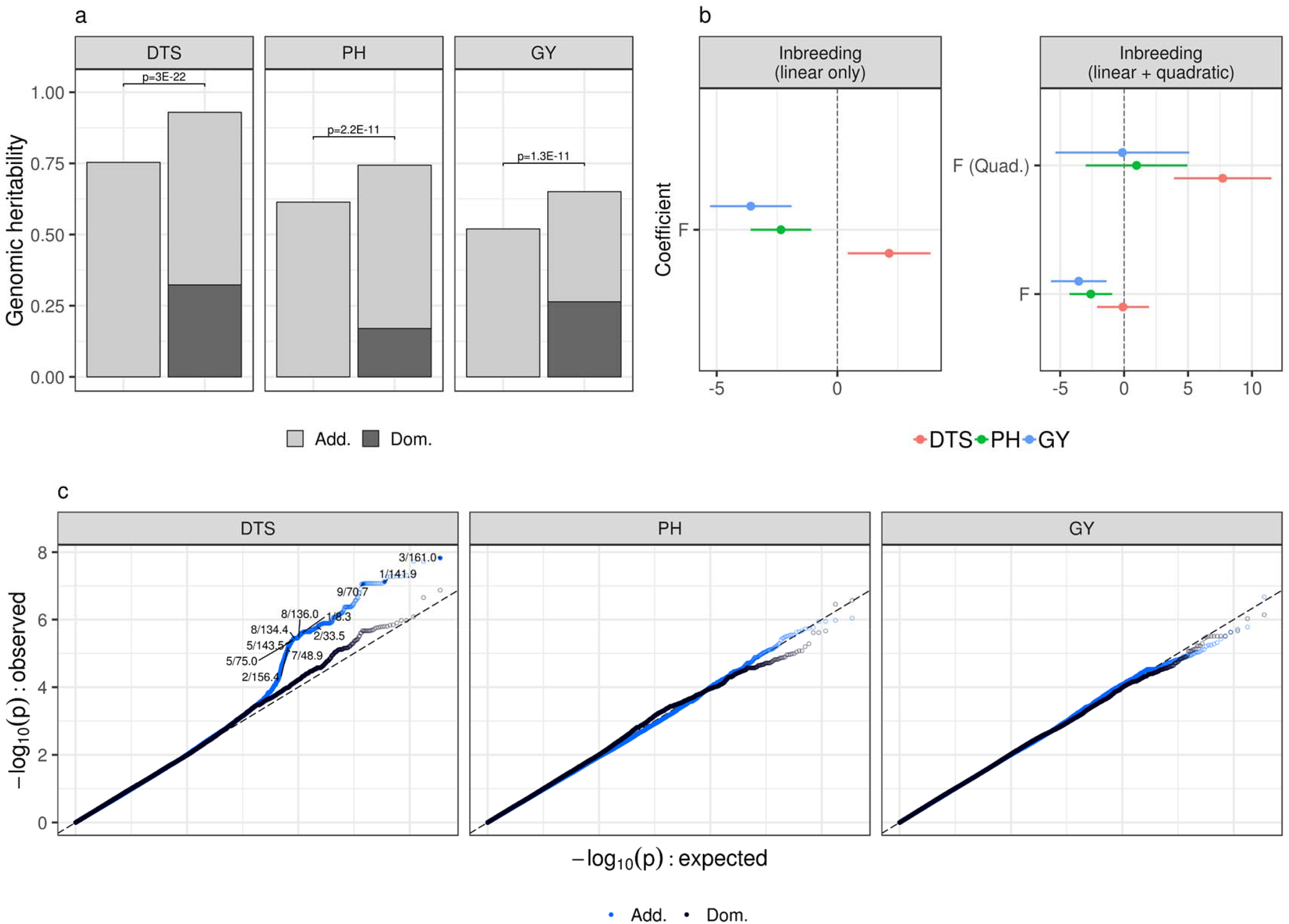
Dominance gene action is a plausible mechanism on hybrid vigor for plant height (PH) and grain yield (GY), but not for days to silking (DTS), in Ames/PHZ51+B47. (a) Partition of variance by additive and dominance effects in genome-wide polygenic models; genomic heritability: proportion of variance among genotype means captured by additive (Add.) or dominance (Dom.) marker effects; p: p-values from likelihood ratio tests. (b) Estimated effects of genomic inbreeding (point and 95% confidence interval). Effects are shown in unit of standard deviations for each trait. F: linear effect; F (Quad.): quadratic effect. (c) Quantile-quantile plot for joint estimates of additive effects (‘Add.’) and dominance effects (‘Dom.’). Effects of SNPs were deemed significant if their false discovery rate (FDR) was lower than 0.05 and if they were not within 1 Mb of SNPs of more significant effects (effects with lower p-values). SNPs with significant effects are designated by chromosome number and genomic position in Mb.

Genomic relationships for epistatic effects were highly correlated with those for additive and/or dominance effects (e.g., *r* > 0.99 between additive and additive×additive relationships). Therefore, we did not assess epistatic effects by partition of genomic variance. Despite this limitation, we further investigated the plausibility of dominance as a genetic mechanism underlying heterosis, by using evidence based on oligogenic effects (QTL effects) and directional effects.

#### Effects of QTL were significant for days to silking but they did not suggest dominance gene action

Effects of QTL were inferred by GWAS models and Bayesian sparse linear mixed models (BSLMMs). Signals from GWAS models and BSLMMs were concordant, and revealed multiple significant QTL effects for DTS (Figure S3). There were five and seven high-confidence QTL (FDR ≤ 0.05 and WPIP ≥ 0.5) for DTS in Ames/PHZ51+B47 and NAM/PHZ51, respectively (Figure S3, Table S1). For PH and GY, no QTL effects were significant except for one QTL for GY in NAM/PHZ51 (Table S1).

GWAS and BSLMM signals for DTS showed limited consistency between Ames/PHZ51+B47 and NAM/PHZ51 (Figure S3), with no overlap of high-confidence QTL across panels (Table S1). This inconsistency could be due to genetic interactions (dominance and/or epistasis), genotype-by-environment interactions, or differential amount of information about SNP effects (different levels of power, due to differences in allele frequency and sample size).

To test whether dominance contributed to QTL effects we conducted a GWAS for additive and dominance QTL effects in Ames/PHZ51+B47. Multiple additive effects appeared significant for DTS, with significant QTL effects (FDR ≤ 0.05) in chromosomes 3, 1, and 9 (Figure 3c). But dominance effects were not significant (FDR > 0.30) (Figure 3c), so factors causing the inconsistency in QTL effects for DTS probably did not involve dominance. Besides, genetic effects did not appear to be sensitive to environments for DTS (Table 2), and there were no systematic differences in allele frequency that could explain difference in significance of QTL across panels (Table S1). Thus, it is plausible that higher-order genetic interactions (epistasis) caused the difference in QTL significance for DTS across panels.

#### Effects of inbreeding pointed to dominance for plant height and grain yield and higher-order genetic interactions for days to silking

Under directional dominance, inbreeding should be linearly related to fitness, but such relationship will tend to be nonlinear under higher-order epistatic interactions such as dominance×dominance interactions (Crow and Kimura 1970). To test whether dominance contributed to genotypic variability by directional effects, we assessed linear and quadratic effects of genomic inbreeding (F) on agronomic traits. For PH and GY in Ames/PHZ51+B47, only linear effects of genomic inbreeding were significant (Figure 3b). Moreover, these effects were on par with their expected impact on fitness, since genomic inbreeding was negatively associated with PH and GY. For DTS in Ames/PHZ51+B47, only the quadratic effect of genomic inbreeding was significant (Figure 3b). Such nonlinear effect implied epistatic gene action for DTS, in the form of SNP×SNP interactions or SNP×background interactions (e.g., differential effects of markers in SS, NSS or semi-tropical genotypes). Along with the lack of dominance QTL effects, the lack of linear effects suggested that dominance is not a predominant genetic mechanism underlying heterosis for DTS.

Despite the high significance of directional effects for all traits in Ames/PHZ51+B47, similar effects were not significant in NAM/PHZ51 (Table S2), possibly because of lower variance and lower range of genomic inbreeding values in this panel (Lynch and Walsh 1998). In fact, variances of *F* and *F*^2^ were respectively 3.9 and 58 times smaller in NAM/PHZ51 (where maximum *F* was only 0.15) compared to Ames/PHZ51+B47 where *F* could be as high as 0.55, for hybrids such as B37×B47 (Table S2).

### Heterosis for all traits may be caused by complementation in proximal gene regions

#### Polygenic effects were enriched in genic regions for all traits

Partition of genomic variance by proximity to annotated genes was significant for all traits in Ames/PHZ51+B47 and NAM/PHZ51, based on likelihood ratio tests combined by Fisher’s method (*p* < 0.01; Table S3; Figure 4a). As suggested by the high correlation in significance (-log_10_(*p*)) between Ames/PHZ51+B47 and NAM/PHZ51 (*r*=0.92), the higher significance of partitions in NAM/PHZ51 could be due to a systematic increase in statistical power, due in part to the larger sample size in NAM/PHZ51 (*n*=1640 vs. *n*=1106).

**Figure 4.**
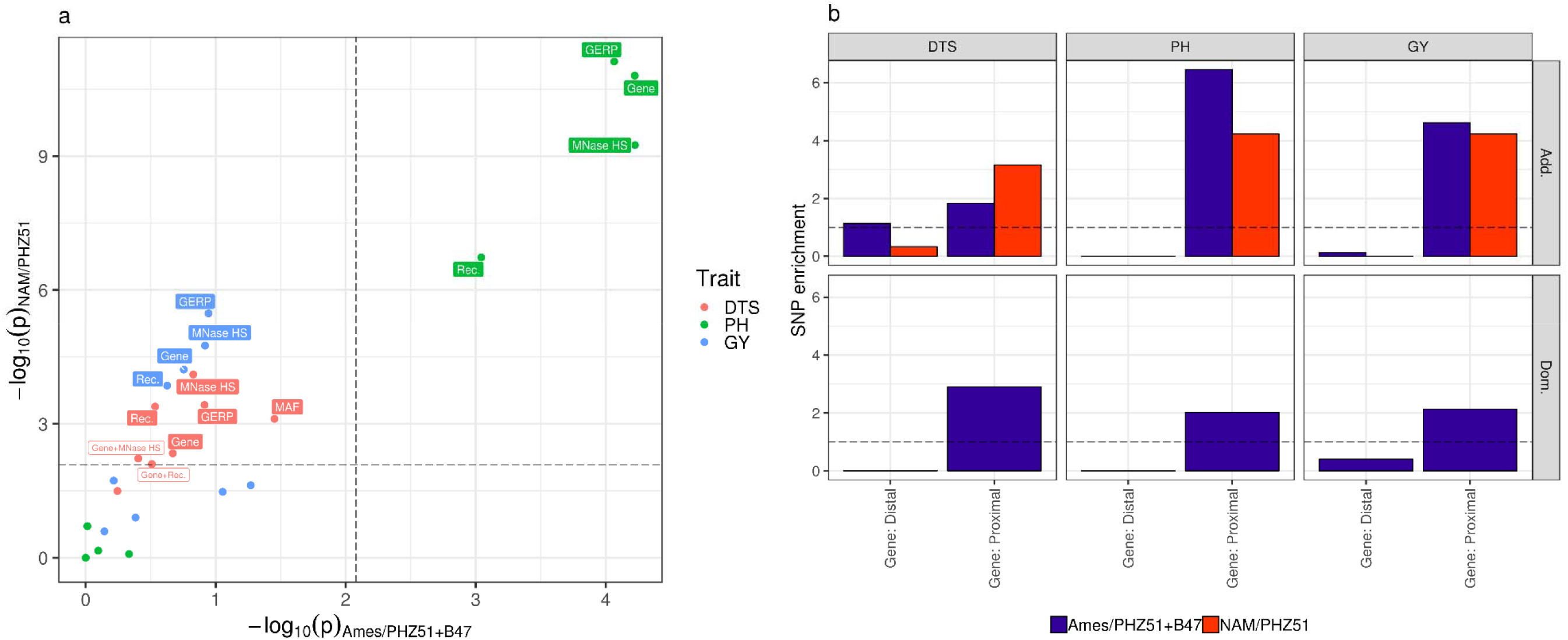
Effects of SNPs on hybrid vigor are enriched in proximal gene regions for days to silking (DTS), plant height (PH) and grain yield (GY), in Ames/PHZ51+B47 and NAM/PHZ51. (a) Significance of variance partition by gene proximity (Gene), structural features (Rec., MNase HS) or evolutionary features (MAF, GERP), and variance partition after accounting for gene proximity (Gene+Rec., Gene+MNase HS, Gene+MAF, Gene+GERP); p-values were obtained by likelihood ratio test comparing the functional model to a baseline model with no partition for the feature of interest (e.g. Gene vs. unpartitioned model, Gene+MAF vs. Gene); dashed lines correspond to thresholds for significance in either panel, after adjustment by Bonferroni correction. Text refers to significant features after Bonferroni correction, based on p-values in either panel (open boxes) or p-values in both panels combined by Fisher’s method (full boxes) (Table S3). (b) Enrichment of SNP heritability, for additive effects (Add.) and dominance effects (Dom.), by bin for gene proximity (Gene). Proximal: ≤ 1 kb of an annotated gene; Distal: ≥ 1 kb from an annotated gene.

Observed SNP enrichments by gene-proximity classes were concordant across panels and traits; they indicated that the magnitude of polygenic effects tended to be higher near genic regions (Figure 4b). Moreover, the proportion of variance explained by gene-proximal SNPs was consistently larger than explained by gene-distal SNPs, except for additive effects in non-genic regions for DTS in Ames/PHZ51+B47 (43% of genomic variance in non-genic regions vs. 22% in genic regions; Table S4). Therefore, there was supporting evidence for genotypic variability arising through genetic effects in genic regions, especially for PH for which enrichment near annotated genes was highly significant in both panels.

#### Enrichment of polygenic effects in low-recombination regions and evolutionarily constrained loci was unclear

Partition of genomic variance explained by recombination rate, chromatin openness, MAF, and GERP scores was significant for DTS in NAM/PHZ51 only (all features), for PH in both panels (all features except MAF), and for GY in NAM/PHZ51 only (all features except MAF) (Figure 4a, Table S3). SNP enrichments in both panels indicated that the magnitudes of polygenic effects tended to be larger at low-diversity loci (low MAF and high GERP scores) and in euchromatic regions (open chromatin and moderate-to-high recombination rates) (Figure S4). However, none of them were significant after accounting for gene proximity, based on likelihood ratio tests combined by Fisher’s method (p > 0.01; Table S3). Because evolutionary constraint and chromatin structure are positively associated with gene density, enrichment at these features may have been due to SNP enrichment by gene proximity.

### Genomic prediction models were improved by dominance effects and functional features

#### Polygenic dominance effects increased prediction accuracy for plant height

Predictions from GBLUP models trained in Ames/PHZ51+B47 were significantly accurate in NAM/PHZ51 (prediction accuracy significantly different from zero) for DTS and PH, but not for GY (Table 3). The absence of predictive ability for GY may be explained by very strong genotype-by-environment interactions (Table 2). Prediction accuracy was highest for DTS, consistently with genomic heritability in Ames/PHZ51+B47 being the highest (Figure 3a) and genetic effects being the most concordant across panels (Table 2). Predictions from DGBLUP models trained in Ames/PHZ51+B47 and tested in NAM/PHZ51 were significantly more accurate than GBLUP for PH (*p* = 0.021), with no significant differences in accuracy for DTS and GY (Table 3). Therefore, accounting for polygenic dominance effects should not be detrimental to genomic prediction models (despite the higher complexity of DGBLUP compared to GBLUP) and may even increase their accuracy.

QTL effects, as estimated from BSLMMs (sparse effects), generated predictions that were significantly accurate in NAM/PHZ51 for DTS (Table S5). Predictions from sparse effects were also significantly accurate for PH, but they actually recapitulated polygenic effects, since their predictions were highly correlated to those from a purely polygenic model (RR-BLUP) (*r* = 0.87 for PH vs. *r* = 0.63 for DTS; Figure S6). Despite QTL effects being highly significant for DTS, a BSLMM simultaneously fitting sparse and polygenic effects did not result in significant gains in prediction accuracy for any trait, compared to RR-BLUP. Therefore, even when QTL effects could capture a significant part of the genotypic variability, they did not seem useful for genomic prediction. Likewise, directional effects (genomic inbreeding) explained a significant part of genotypic variability in Ames/PHZ51+B47 (Figure 3b, Table S2), but incorporating them into DGBLUP models resulted in small and non-significant differences in prediction accuracy in NAM/PHZ51 populations (Table S6).

#### Partition of polygenic effects by gene proximity increased prediction accuracy for all traits

The amplitude of polygenic effects was inflated near annotated genes in both panels for all traits (Figure 4b). Accordingly, partitioning genomic variance by gene proximity increased prediction accuracy for DTS (p = 3.3×10^-4^), PH (p = 0.023), and GY (p = 0.085), compared to a DGBLUP model (Table 4).

Other features than gene proximity also resulted in significant gains in prediction accuracy. Partitioning genomic variance by recombination-rate classes increased prediction accuracy for GY (Table 4); besides, recombination rate and GERP scores contributed to improvements that were significant, but only when enrichment by gene proximity was omitted from prediction models for DTS (Table 4). Even when classes based on MNase HS and GERP scores did not yield significant gains in prediction accuracy, the high SNP enrichments achieved by these features (~32-fold for open-chromatin regions and ~8-fold for GERP > 0) could be of practical interest for SNP selection in prediction analyses (Figure S4, Table 4). Therefore, although gene proximity appeared most useful and meaningful in our study, the value of structural and evolutionary features for genomic prediction would deserve further investigation.

**Table 3.** Prediction accuracy in NAM/PHZ51 by polygenic dominance effects in Ames/PHZ51+B47

| Trait | Prediction accuracy (p-value) | Difference in prediction accuracy (p-value) |
| --- | --- | --- |
|  | GBLUP | Dominance GBLUP |
| **DTS** | 0.331 (< 0.001) | -0.013 (0.21) |
| **PH** | 0.235 (< 0.001) | +0.023 (0.021) |
| **GY** | -0.001 (0.96) | -0.010 (0.33) |

Prediction accuracy: average correlation between observed and predicted phenotypes in NAM/PHZ51 over 24 populations by the GBLUP model; Difference in prediction accuracy: difference between the dominance GBLUP (DGBLUP) model and the GBLUP model. Significance of average prediction accuracies (non-zero mean) and estimated differences in prediction accuracy (non-zero difference) was assessed by t-tests paired by NAM population.

**Table 4.**
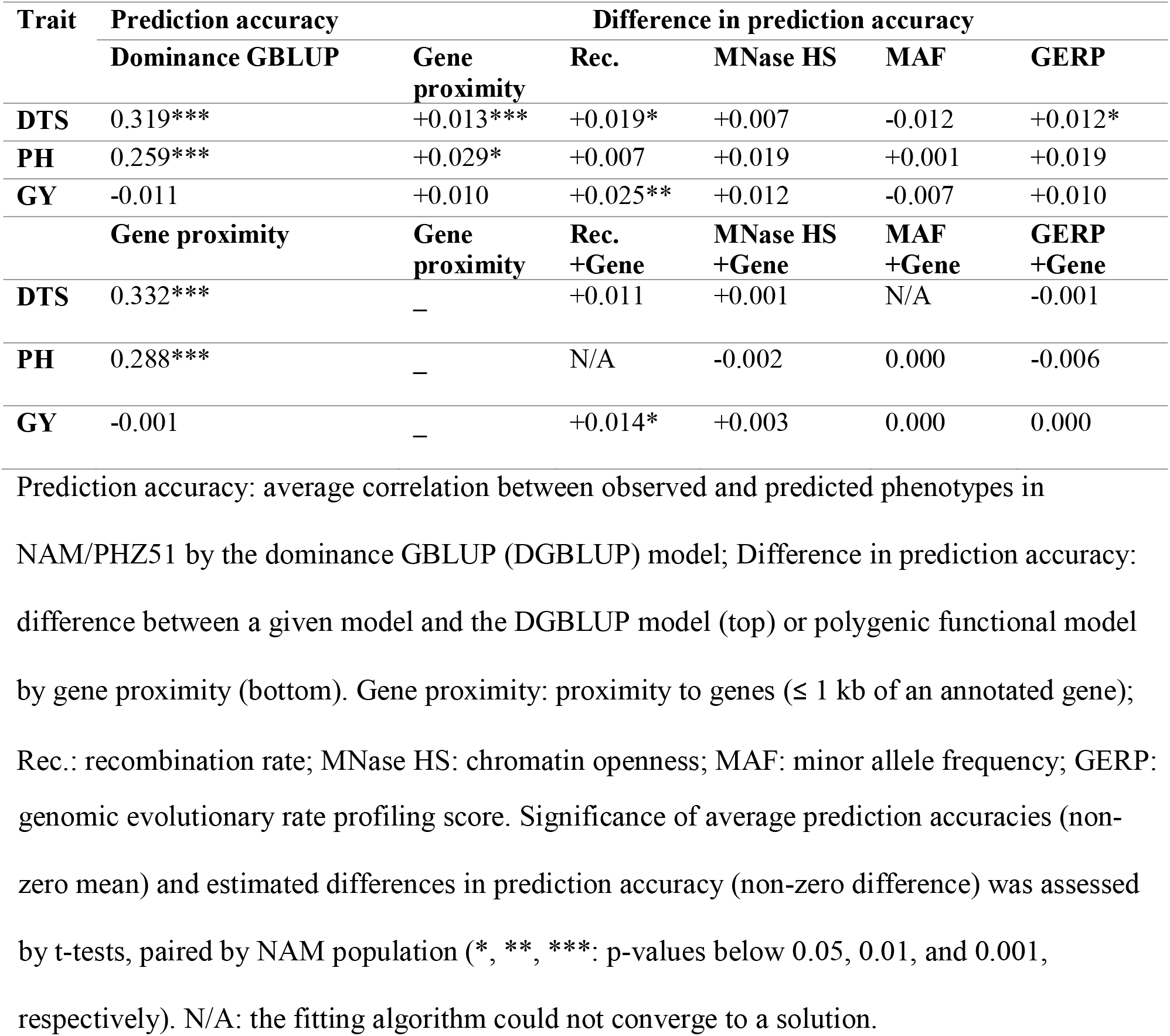
Prediction accuracy in NAM/PHZ51 by functional features in Ames/PHZ51+B47

## DISCUSSION

### Do additive and dominance gene actions adequately capture true genetic effects?

All traits displayed a significant proportion of variance explained by dominance effects (Figure 3a). However for DTS, there was conflicting evidence about the importance of dominance: (i) no significant dominance QTL effects despite significant additive QTL effects (Figure 3c) and (ii) significant quadratic effects of genomic inbreeding, without any linear effect (Figure 3b). Such evidence indicates that DTS should probably be analyzed under more complex genetic models involving epistatic interactions, possibly reflecting the complex molecular pathways underlying flowering time (e.g., photoperiod genes; Yang et al. 2013, Blümel et al. 2015, Minow et al. 2018). In this study, genomic variance in Ames/PHZ51+B47 could not be partitioned reliably by additive, dominance, and epistatic effects. Indeed, genomic relationships for pairwise epistatic effects were highly correlated with those for additive effects. Moreover, epistatic effects in linear mixed models vary depending on how marker variables are centered, in a way that can be arbitrary (Martini et al. 2016, Martini et al. 2017). However, further analyses to investigate the contribution of epistatic effects to genomic variance is merited (Jiang and Reif 2015). Investigating epistatic effects would probably require large panels with more testers (male parents), but also efficient methodologies to restrict the number of interactions (e.g., only interactions between homeologs; Santantonio et al. 2018) and the types of effects involved (e.g., only SNP×SNP interactions such as additive×additive effects, or SNP×background interactions such as SNP×PC effects; Ramstein et al. 2018).

For PH and GY, there was concordant evidence for prevalent dominance effects: (i) significant variance partition by dominance effects (Figure 3a) and (ii) a significant linear effect of genomic inbreeding, without any quadratic effect (Figure 3b). Therefore, additive and dominance effects may parsimoniously capture genetic effects for PH and GY. These results contrast with those from previous studies on hybrid maize, which showed low contribution of non-additive genetic effects to genotypic variability. Critically, those studies were based on panels derived solely from crosses between different heterotic groups, e.g., Flint×Dent (Technow et al. 2014, Giraud et al. 2017) or SS×NSS (Kadam et al. 2016). Therefore, complementation effects were relatively consistent across hybrids, such that variability for specific combining ability (contributed by dominance and/or epistasis) was low. In contrast, one of our panels (Ames/PHZ51+B47) showed strong variation for complementation effects, because it represents a variety of genetic contexts (SS×NSS, SS×SS, Semi-tropical×SS, etc.). Therefore, it was better-suited to represent the differential levels of complementation effects in maize and reveal the importance of dominance across maize hybrids.

### What is the biological basis for enrichment of SNP effects by gene proximity?

Analyses of SNP enrichment pointed to genetic effects arising mostly from genic regions (proximal SNPs, ≤ 1 kb from annotated genes). The relevance of genic regions for depicting hybrid vigor is consistent with hypotheses about biological causes of heterosis related to gene expression, namely, (i) non-additive inheritance of gene expression and (ii) nonlinear effects of gene expression on agronomic traits (Springer and Stupar 2007, Schnable and Springer 2013). In maize, studies have generally reported that most genes have an additive mode of inheritance for expression levels (e.g., Swanson-Wagner et al. 2006, Stupar and Springer 2006, Zhou et al. 2018), with proportions of non-additive gene actions ranging from ~10% (Paschold et al. 2012) to ~35% (Marcon et al. 2017). Proposed mechanisms for non-additive gene expression include complementation with respect to regulatory motifs or transcription factors, and presence/absence variation (Paschold et al. 2012, Marcon et al. 2017, Zhou et al. 2018). Importantly, studies on gene expression in maize have also suggested that non-additive gene expression may not account entirely for heterosis (Swanson-Wagner et al. 2006, Stupar and Springer 2006). Therefore, the genome-wide patterns of apparent dominance at gene regions observed here (Figure 4b) might have also emerged from nonlinear effects of gene expression on agronomic traits. Evidence for this type of effects in maize include intermediate gene expression harboring minimal burden of deleterious mutations in diverse maize inbred lines (Kremling et al. 2018) and biological results in support for the gene balance hypothesis (Birchler and Veitia 2010), which postulates that genes that are highly connected (in pathways, protein complexes, etc.) should be expressed in relative amounts under a stoichiometric optimum (Birchler et al. 2001). Optimal expression levels under gene balance constitute nonlinear effects of gene expression, and may contribute to non-additive genetic effects (Birchler and Veitia 2010). Ideally, future research about the biological basis for enrichment of SNP effects in genic regions will involve gene expression data in diverse hybrid panels, and will shed light onto the relative importance of such phenomena on heterosis.

The hybrid panels under assay were relatively large, so that we could gain useful insight about SNP enrichments by functional classes. However, some biological and genetic hypotheses could not be tested due to limited power and resolution in our analyses. For example, we could not account for enrichment in genic regions and concurrently assess the functional importance of evolutionary constraint or repulsion phase linkage. Moreover, because of high correlation between genomic relationships from different bins (r > 0.99), we did not consider finer partitions for different levels of deleteriousness (more than two GERP-score classes; Wang et al. 2017) or relevance of chromatin openness at different tissues (root and/or shoot; Rodgers-Melnick et al. 2016). Therefore, larger sample sizes will be critical to investigate finer partitions in functional models, allowing higher resolution and better control of confounding factors like gene density.

### Are enrichments in genic regions and dominance effects useful for genomic selection?

The practical relevance of dominance effects and SNP enrichments were evaluated here by genomic prediction in each NAM/PHZ51 population, based on models trained in a different panel (Ames/PHZ51+B47). Therefore, prediction models were assessed for their ability to sustain accuracy across distinct population backgrounds. Enrichment of SNP effects increase the representation of loci that are more likely to be causal; therefore, enrichment procedures like QTL detection or variance partition can improve the accuracy of genomic prediction models. However, as genetic effects vary from one population background to another, enrichments about small functional classes (e.g., a few GWAS hits) lose their potential. This caveat was exemplified by differences in QTL effects for DTS between Ames/PHZ51+B47 and NAM/PHZ51, and the consequent lack of gain in accuracy by prediction models based on QTL effects (BSLMMs) (Table S5). Similarly, Spindel et al. (2016) showed benefits of major QTL effects for prediction of flowering time in rice, but only when QTL were detected on the target breeding populations. Contrary to enrichments about QTL, enrichments about larger functional classes (e.g., gene-proximal SNPs) should result in gains of prediction accuracy that are robust to differences in population backgrounds and consistent over traits, as was observed here (Table 4). Likewise, Gao et al. (2017) reported gains in genomic prediction accuracy by prioritizing genic SNPs, in mouse, drosophila, and rice (increases in predictive ability averaging +0.013, similar to those realized in this study). Therefore, gains in prediction accuracy by gene proximity should be expected under a broad range of population and species contexts.

While SNP enrichments by gene proximity appeared beneficial for all traits, incorporating dominance resulted in gains in prediction accuracy for PH only (Table 3). The absence of gain in prediction accuracy for DTS and GY illustrates possible reasons for disagreement between quality of fit and prediction accuracy often observed in genomic prediction studies. For DTS, incorporating dominance effects resulted in *statistically* significant improvements in fit, but a genetic model accounting for epistatic interactions appeared more plausible according to analyses of QTL and genomic inbreeding. Therefore, the choice of the prediction procedure should probably come from multiple pieces of evidence in favor of a given genetic model, rather than a single statistical test about the prediction model. In the case of GY, a genetic model based only on additive and dominance effects seemed plausible in Ames/PHZ51+B47, but the dependency of these effects on environmental backgrounds hindered predictions in NAM/PHZ51. Therefore, prediction of hybrid performance for GY should probably accommodate genotype-by-environment interactions, through models based on environmental covariates related to temperature, radiation, or soil water potential (Li et al. 2018, Millet et al. 2019).

## Conclusions

Our analyses point to genetic models in hybrid maize which involve interactive effects and emphasize genic regions. While dominance may be relevant to all three traits, epistasis seemed particularly important for DTS, and interactions with environments seemed critical for GY. Consequently, genomic prediction models were improved by dominance effects for PH only, while they benefited from SNP enrichment in genic regions for all traits. These results call for further investigation about the biological basis of genetic complementation, and the underlying interactive effects which could enable more robust prediction of hybrid vigor.

## ACKNOWLEDGEMENTS

This work was funded by NSF Plant Genome Program (IOS 0820619 and 1238014) and USDA-ARS. Graduate work of SJL, work of ESE, and IA10 trials were partially funded by Syngenta.

