## Supplementary information for "The relevance of dominance and functional annotations to predict agronomic traits in hybrid maize"

**Table S1 – High-confidence QTL effects in Ames/PHZ51+B47 and NAM/PHZ51, based on GWAS models and BSLMMs**

| Trait | Chr. | Position | Ames/PHZ51+B47 | | | | NAM/PHZ51 | | | |
| --- | --- | --- | --- | --- | --- | --- | --- | --- | --- | --- |
|  |  |  | **AF** | **Effect** | **p (FDR)** | **WPIP** | **AF** | **Effect** | **p (FDR)** | **WPIP** |
| DTS | 1 | 5,819,499 | 0.76 | 0.14 | 0.37 (0.99) | < 0.01 | **0.86** | **0.76** | **1.8×10^-10^ (8×10^-7^)** | **0.93** |
|  | 1 | 6,246,295 | 0.61 | -0.31 | 0.022 (0.81) | < 0.01 | **0.80** | **0.48** | **1.1×10^-5^ (0.011)** | **0.92** |
|  | 2 | 225,200,515 | 0.99 | 1.20 | 0.019 (0.80) | < 0.01 | **0.91** | **0.74** | **4.1×10^-7^ (7.4×10^-4^)** | **0.73** |
|  | 2 | 225,463,382 | 0.61 | 0.18 | 0.19 (0.99) | < 0.01 | **0.77** | **0.59** | **2.2×10^-8^ (6.2×10^-5^)** | **0.90** |
|  | 3 | 161,016,441 | **0.58** | **-0.86** | **7.3×10^-9^ (2.6×10^-3^)** | **0.55** | 0.88 | -0.51 | 2.8×10^-4^ (0.13) | 0.01 |
|  | 4 | 202,429,351 | **0.96** | **1.74** | **1.9×10^-7^ (0.014)** | **0.51** | 0.89 | 0.03 | 0.86 (1.00) | < 0.01 |
|  | 7 | 110,855,373 | **0.99** | **3.69** | **6.3×10^-10^ (4.4×10^-4^)** | **0.94** | 0.95 | 0.07 | 0.75 (1.00) | 0.01 |
|  | 8 | 126,761,299 | **0.90** | **-1.04** | **3.8×10^-7^ (0.018)** | **0.59** | 0.97 | -0.70 | 0.023 (0.90) | 0.01 |
|  | 9 | 128,205,378 | 0.67 | 0.19 | 0.21 (0.99) | < 0.01 | **0.74** | **0.55** | **2.3×10^-6^ (3.0×10^-3^)** | **0.89** |
|  | 9 | 146,679,383 | **0.81** | **-0.73** | **3.5×10^-6^ (0.037)** | **0.72** | 0.95 | -0.54 | 0.010 (0.72) | < 0.01 |
|  | 10 | 94,059,383 | 0.96 | 0.05 | 0.88 (1.00) | < 0.01 | **0.95** | **2.23** | **1.3×10^-25^ (4.2×10^-20^)** | **0.99** |
|  | 10 | 94,453,179 | 0.97 | 0.80 | 0.041 (0.88) | < 0.01 | **0.93** | **2.12** | **9.6×10^-31^ (2.4×10^-24^)** | **1.01** |
| GY | 7 | 175,528,863 | 0.79 | -0.05 | 0.54 (0.99) | 0.04 | **0.97** | **-1.09** | **2.0×10^-8^ (0.013)** | **0.70** |

Chr.: Chromosome; Position: physical location; AF: frequency of major allele (as per the Hapmap 3.2.1 panel); Effect: estimate of SNP effect from GWAS; p (FDR): p-value by Wald tests and false discovery rate estimated by the method of Benjamini and Hochberg (1995); WPIP: posterior inclusion probability for non-zero SNP effects in 500-kb sliding windows (250-kb steps), based on BSLMM (Guan and Stephens 2011). High-confidence QTL effects are characterized by FDR ≤ 0.05 and WPIP ≥ 0.5.

**Table S2 ­– Summary statistics about genomic inbreeding in each set and the significance of its effects, by trait**

| Set | Variance | Range | Trait | p |
| --- | --- | --- | --- | --- |
| Ames/  PHZ51+B47 | *F*: 3.4×10^-3^  *F*^2^: 2.2×10^-4^ | *F*: (-0.091, 0.55)  *F*^2^: (2.2×10^-9^, 0.30) | DTS | 1.8×10^-5^ |
|  |  |  | PH | 1.3×10^-3^ |
|  |  |  | GY | 1.8×10^-4^ |
| NAM/  PHZ51 | *F*: 8.8×10^-4^ *F*^2^: 3.8×10^-6^ | *F*: (-0.093, 0.15) *F*^2^: (5.3×10^-8^, 0.023) | DTS | 0.53 |
|  |  |  | PH | 0.32 |
|  |  |  | GY | 0.14 |

Effects tested were *F* and *F*^2^, where *F* is the genomic inbreeding coefficient; p-values were obtained from Wald tests for effects of *F* and *F*^2^ simultaneously.

**Table S3 – Significance of variance partition by panel, by functional feature**

| Set | Trait | p-value for variance partition | | | | |
| --- | --- | --- | --- | --- | --- | --- |
|  |  | **Gene** | **Rec.** | **MNase HS** | **MAF** | **GERP** |
| Ames/  PHZ51+B47 | DTS | 0.21 | 0.29 | 0.15 | 0.035 | 0.12 |
|  | PH | 6.0×10^-5^ | 9.1×10^-4^ | 6.0×10^-5^ | 0.80 | 8.6×10^-5^ |
|  | GY | 0.18 | 0.24 | 0.12 | 0.088 | 0.11 |
| NAM/  PHZ51 | DTS | 4.6×10^-3^ | 4.1×10^-4^ | 7.8×10^-5^ | 7.7×10^-4^ | 3.8×10^-4^ |
|  | PH | 1.6×10^-11^ | 1.9×10^-7^ | 5.6×10^-10^ | 0.70 | 7.5×10^-12^ |
|  | GY | 6.1×10^-5^ | 1.4×10^-4^ | 1.8×10^-5^ | 0.033 | 3.4×10^-6^ |
| Combined  (Fisher’s method) | DTS | 7.8×10^-3^ | 1.2×10^-3^ | 1.4×10^-4^ | 3.1×10^-4^ | 5.1×10^-4^ |
|  | PH | 3.3×10^-14^ | 4.0×10^-9^ | 1.1×10^-12^ | 0.88 | 2.3×10^-14^ |
|  | GY | 1.3×10^-4^ | 3.7×10^-4^ | 3.0×10^-5^ | 0.020 | 6.0×10^-6^ |
|  |  | **Gene** | **Gene+**  **Rec.** | **Gene+**  **MNase HS** | **Gene+**  **MAF** | **Gene+**  **GERP** |
| Ames/  PHZ51+B47 | DTS | _ | 0.31 | 0.39 | N/A | 0.57 |
|  | PH | _ | N/A | 0.46 | 1.0 | 0.97 |
|  | GY | _ | 0.72 | 0.41 | 0.054 | 0.61 |
| NAM/  PHZ51 | DTS | _ | 8.0×10^-3^ | 6.0×10^-3^ | 1.0×10^-4^ | 0.032 |
|  | PH | _ | 1.0 | 0.83 | 1.0 | 0.20 |
|  | GY | _ | 0.26 | 0.13 | 0.024 | 0.019 |
| Combined  (Fisher’s method) | DTS | _ | 0.017 | 0.017 | N/A | 0.091 |
|  | PH | _ | N/A | 0.75 | 1.0 | 0.51 |
|  | GY | _ | 0.50 | 0.21 | 9.8×10^-3^ | 0.063 |

Significance by gene proximity (Gene), structural features (Rec., MNase HS), and evolutionary features (MAF, GERP); p­-values were obtained by likelihood ratio test comparing the variance component model (estimating variance of additive and dominance marker effects by bin) to the baseline model: no partition (top) or variance partition by gene proximity (bottom). N/A: the fitting algorithm could not converge to a solution.

**Table S4 – Genomic variance captured by functional classes**

| Feature | Bin | Genomic variance  (Proportion of genomic heritability explained) | | | | | |
| --- | --- | --- | --- | --- | --- | --- | --- |
|  |  | **Ames/PHZ51+B47** | | | **NAM/PHZ51** | | |
|  |  | **DTS** | **PH** | **GY** | **DTS** | **PH** | **GY** |
| Gene  proximity | Non-genic | **0.40 (43%)** | 0.00 (0%) | 0.03 (5%) | **0.19 (25%)** | 0.00 (0%) | 0.00 (0%) |
|  | Genic | **0.20 (22%)** | **0.58 (76%)** | **0.36 (55%)** | **0.56 (75%)** | **0.72 (100%)** | **0.49 (100%)** |
|  | Non-genic (D) | 0.01 (1%) | 0.00 (0%) | 0.10 (16%) |  |  |  |
|  | Genic (D) | **0.32 (34%)** | **0.18 (24%)** | **0.16 (25%)** |  |  |  |
| Rec. | < 0.45 | 0.11 (12%) | 0.00 (0%) | 0.01 (2%) | 0.04 (5%) | 0.03 (5%) | 0.01 (2%) |
|  | (0.45, 1.65) | **0.41 (44%)** | **0.17 (22%)** | **0.14 (21%)** | **0.42 (55%)** | **0.22 (31%)** | **0.13 (27%)** |
|  | > 1.65 | 0.08 (9%) | **0.40 (53%)** | **0.23 (35%)** | **0.30 (39%)** | **0.46 (64%)** | **0.35 (71%)** |
|  | < 0.45 (D) | 0.07 (8%) | 0.02 (3%) | 0.06 (9%) |  |  |  |
|  | (0.45, 1.65) (D) | 0.10 (11%) | 0.02 (3%) | 0.07 (11%) |  |  |  |
|  | > 1.65 (D) | **0.15 (16%)** | **0.15 (19%)** | **0.14 (22%)** |  |  |  |
| MNase HS | Dense | **0.17 (18%)** | 0.00 (0%) | **0.27 (42%)** | 0.06 (7%) | 0.00 (0%) | 0.00 (0%) |
|  | Open | **0.43 (46%)** | **0.58 (75%)** | **0.12 (19%)** | **0.70 (93%)** | **0.71 (100%)** | **0.51 (100%)** |
|  | Dense (D) | **0.32 (35%)** | 0.00 (0%) | 0.00 (0%) |  |  |  |
|  | Open (D) | 0.01 (1%) | **0.19 (25%)** | **0.26 (40%)** |  |  |  |
| MAF | < 0.01 | 0.00 (0%) | 0.00 (0%) | 0.00 (0%) | 0.00 (0%) | 0.05 (7%) | **0.17 (36%)** |
|  | (0.01, 0.05) | **0.46 (50%)** | 0.07 (10%) | 0.09 (14%) | **0.34 (46%)** | 0.00 (0%) | 0.00 (0%) |
|  | > 0.05 | **0.14 (15%)** | **0.40 (54%)** | 0.06 (9%) | **0.40 (54%)** | **0.64 (93%)** | **0.31 (64%)** |
|  | < 0.01 (D) | 0.00 (0%) | 0.00 (0%) | 0.00 (0%) |  |  |  |
|  | (0.01, 0.05) (D) | 0.00 (0%) | **0.13 (18%)** | **0.36 (53%)** |  |  |  |
|  | > 0.05 (D) | **0.32 (35%)** | **0.14 (19%)** | **0.16 (24%)** |  |  |  |
| GERP | 0 | **0.46 (49%)** | 0.00 (0%) | **0.16 (24%)** | 0.15 (19%) | 0.00 (0%) | 0.00 (0%) |
|  | > 0 | 0.14 (15%) | **0.58 (76%)** | **0.24 (36%)** | **0.61 (81%)** | **0.72 (100%)** | **0.51 (100%)** |
|  | 0 (D) | 0.00 (0%) | 0.01 (1%) | 0.00 (0%) |  |  |  |
|  | > 0 (D) | **0.32 (35%)** | **0.17 (23%)** | **0.26 (40%)** |  |  |  |

Variance partition by functional features. Gene proximity: proximity to genes (≤ 1 kb); Rec.: recombination rate; MNase HS: chromatin openness; MAF: minor allele frequency; GERP: genomic evolutionary rate profiling score. Genomic variance: variance component at a given bin from polygenic functional model. Proportion of genomic heritability explained: ratio of genomic variance at a given bin over the sum of genomic variances across bins.

**Table S5 – Prediction accuracy in NAM/PHZ51 from oligogenic models fitted in Ames/PHZ51+B47**

| Trait | Prediction accuracy | | Difference in prediction accuracy |
| --- | --- | --- | --- |
|  | **Sparse** | **Polygenic**  **(RR-BLUP)** | **Sparse + polygenic** |
| DTS | 0.180 *** | 0.334 *** | -0.059 |
| PH | 0.150 *** | 0.234 *** | -0.007 |
| GY | 0.016 | 0.002 | +0.003 |

Prediction accuracy: average correlation between observed and predicted phenotypes in NAM/PHZ51 over 24 populations; Polygenic: polygenic effects from RR-BLUP; Sparse: sparse effects from BSLMM, after regressing out polygenic effects from BSLMM; Sparse + polygenic: effect of both sparse and polygenic effects from BSLMM. Significance of average prediction accuracies (non-zero mean) and estimated differences in prediction accuracy (non-zero difference) was assessed by t-tests, paired by NAM population (*, **, ***: p-values below 0.05, 0.01, and 0.001, respectively).

**Table S6 – Prediction accuracy in NAM/PHZ51 by directional effects of inbreeding in Ames/PHZ51+B47**

| Trait | Prediction accuracy | Difference in prediction accuracy | |
| --- | --- | --- | --- |
|  | **DGBLUP** | **Linear only** | **Linear + quadratic** |
| DTS | 0.319 *** | +0.001 | -0.002 |
| PH | 0.259 *** | +0.001 | +0.002 |
| GY | -0.011 | 0.000 | 0.000 |

Prediction accuracy: average correlation between observed and predicted phenotypes in NAM/PHZ51 over 24 populations; Difference in prediction accuracy: difference between a given model and the dominance GBLUP model (DGBLUP); Linear only: DGBLUP incorporating *F* as fixed effect; Linear + quadratic: DGBLUP incorporating *F* and *F*^2^ as fixed effects; *F*: genomic inbreeding coefficient. Significance of average prediction accuracies (non-zero mean) and estimated differences in prediction accuracy (non-zero difference) was assessed by t-tests, paired by NAM population (*, **, ***: p-values below 0.05, 0.01, and 0.001, respectively).

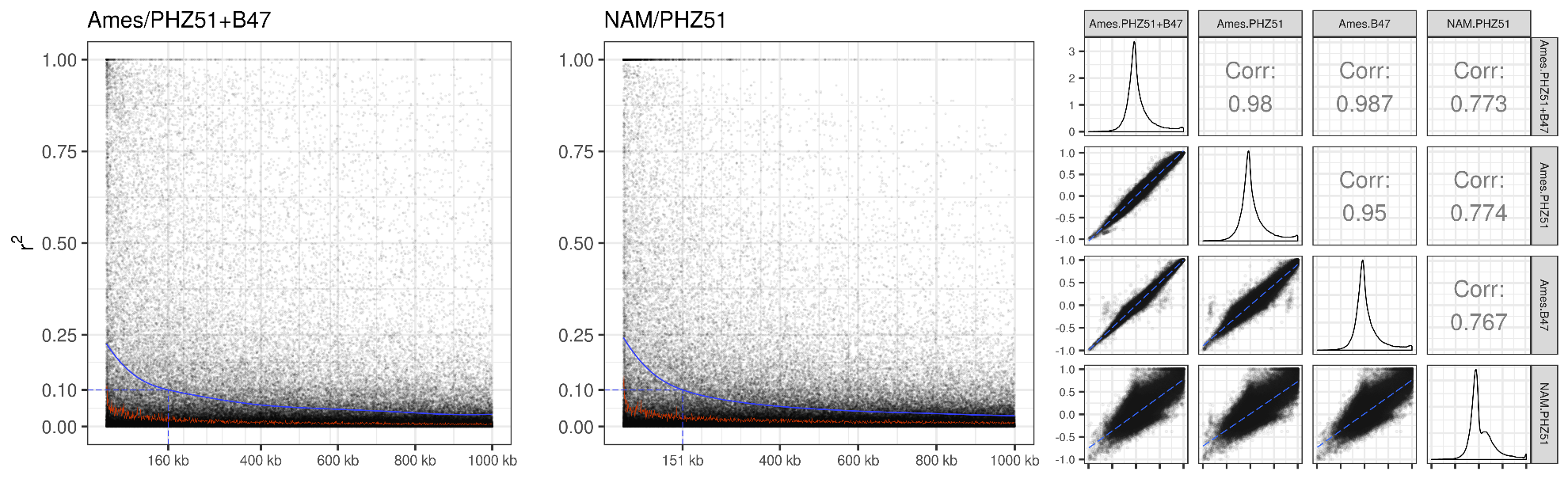
**Figure S1 –** Linkage disequilibrium (LD) in Ames sets (Ames/PHZ51, Ames/B47, Ames/PHZ51+B47) and NAM/PHZ51. Left and middle: decay of LD values (squared correlation of marker scores at markers in pairs, adjusted for population structure and relatedness), as in Mangin *et al.* (2012), in Ames/PHZ51+B47 and NAM/PHZ51 respectively. Right: correlation and scatter plots for adjusted LD values, for each pair of sets. LD values were calculated from random SNP pairs, without sharing of SNPs across pairs.

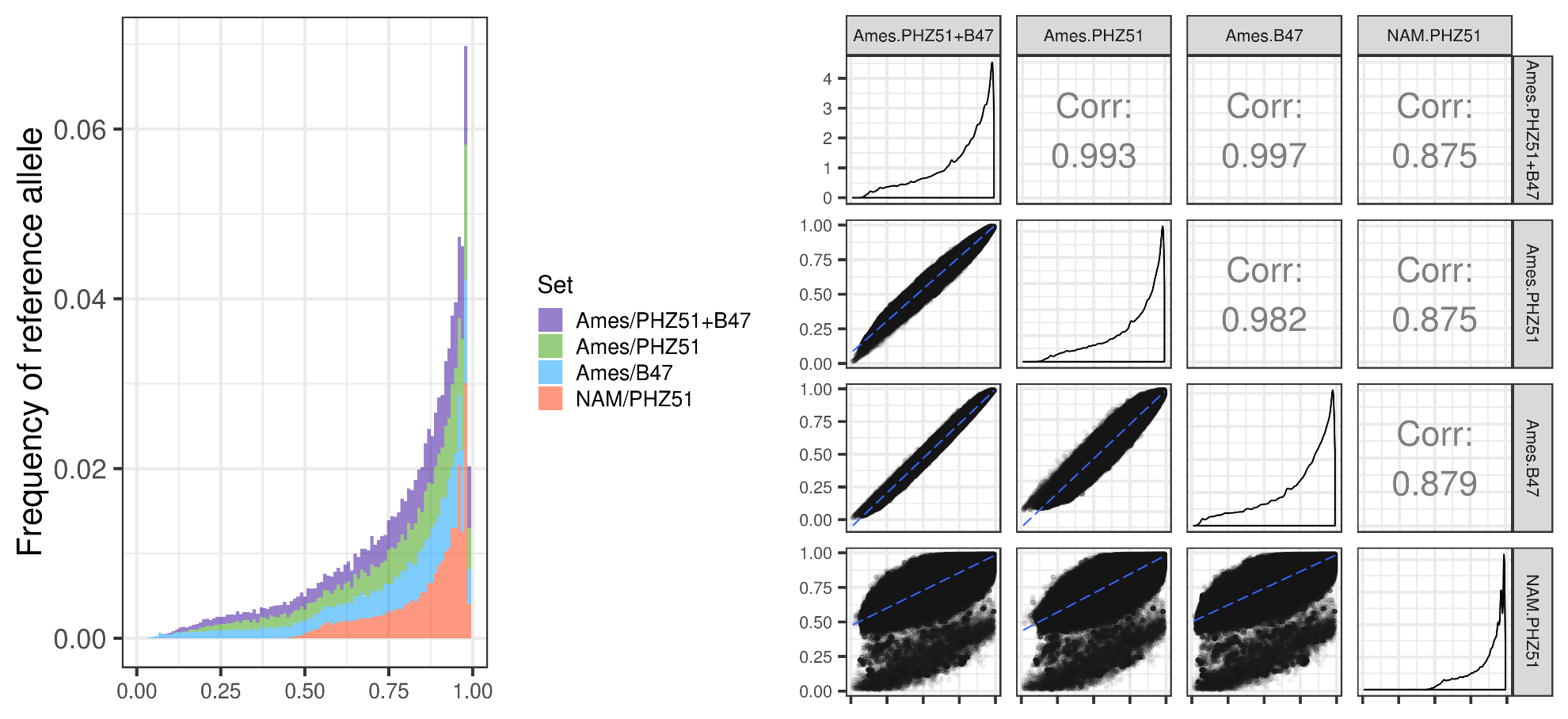
**Figure S2 –** Allele frequency (frequency of reference B73 allele) in Ames sets (Ames/PHZ51, Ames/B47, Ames/PHZ51+B47) and NAM/PHZ51. Left: distribution of allele frequency in each set. Right: correlation and scatter plots for allele frequencies, for each pair of sets.

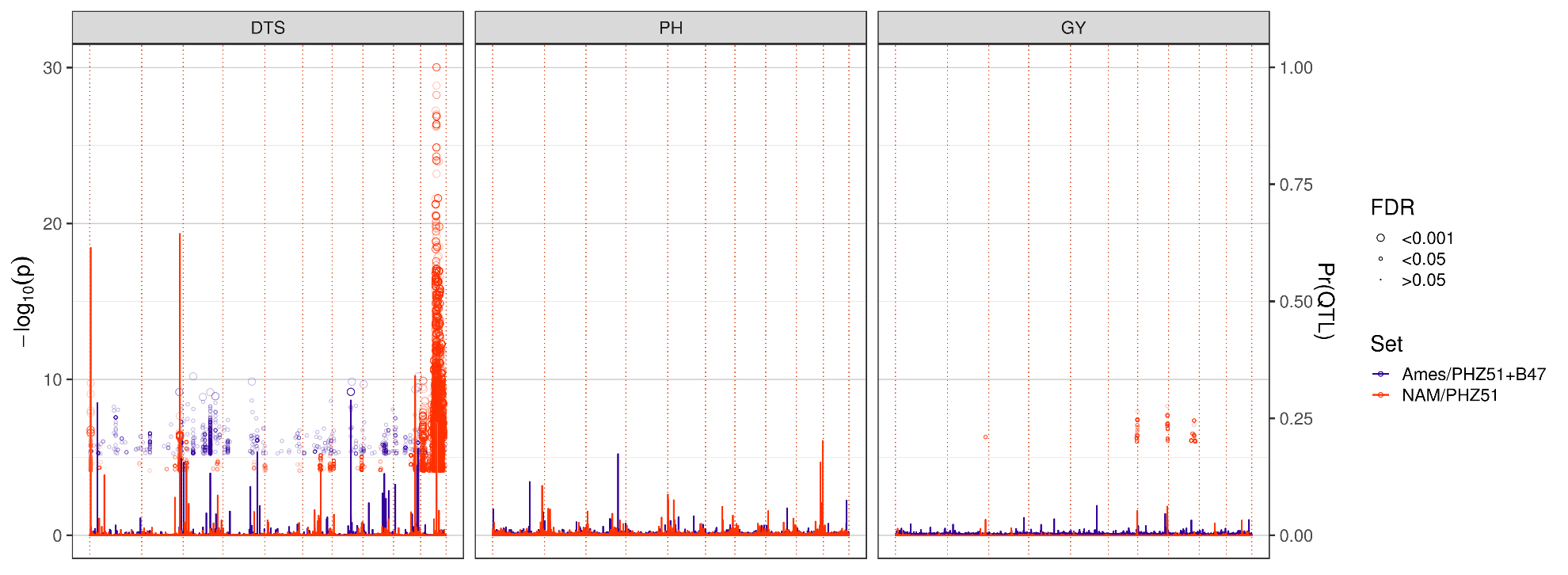

**Figure S3 –** Manhattan plot and BSLMM plot for marginal additive effects. -log_10_(p): significance of SNP effect in GWAS models; p: p-value from Wald tests; significance is only shown for SNPs with FDR < 0.05, with increasing size for higher significance. Pr(QTL): probability of a non-zero sparse effect from BSLMM ($\gamma_{j}$).

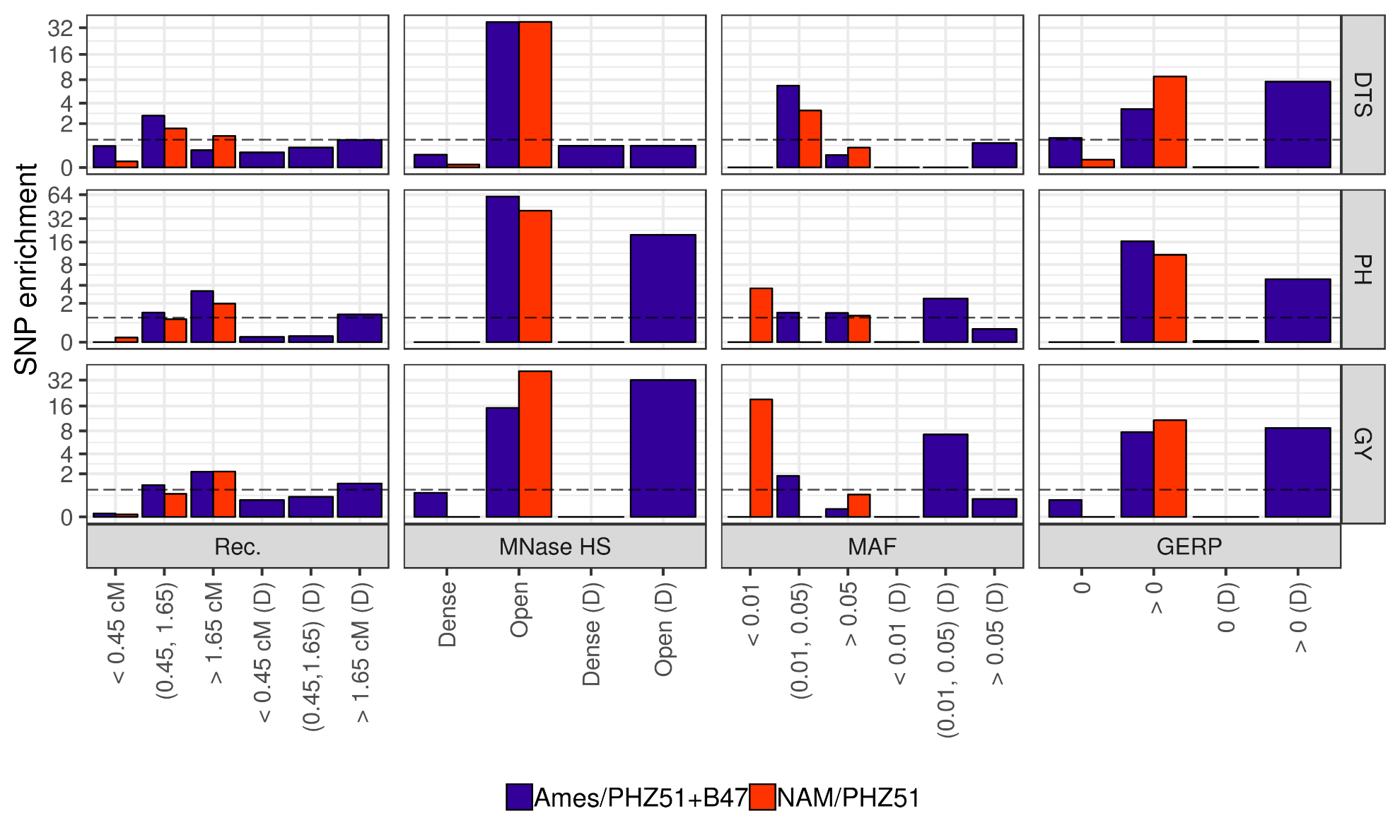

**Figure S4 –** Enrichment of SNP heritability, for additive effects and dominance effects (D), by bin for structural and evolutionary features in Ames/PHZ51+B47 and NAM/PHZ51. Rec.: recombination rate; MNase HS: chromatin openness; MAF: minor allele frequency; GERP: genomic evolutionary rate profiling score.

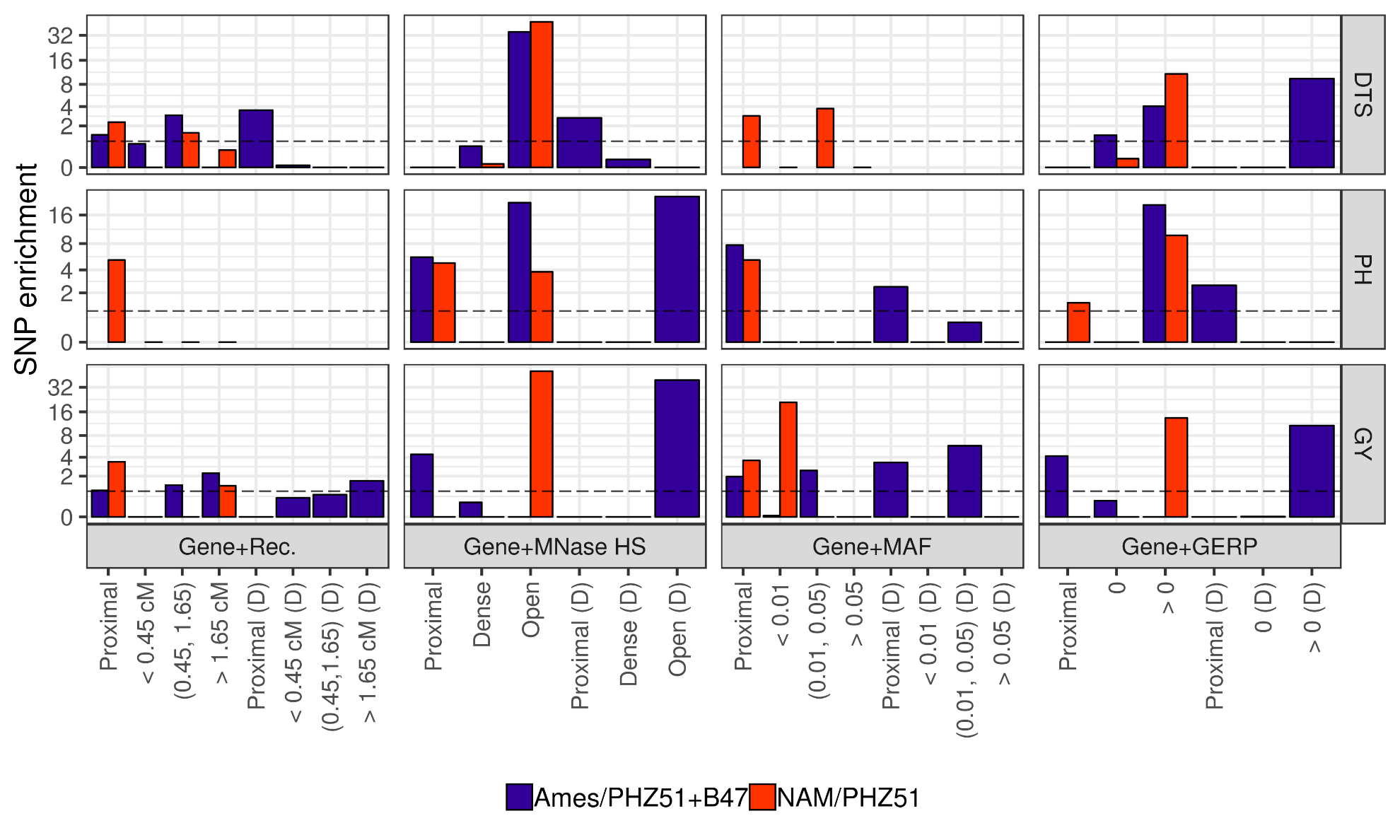

**Figure S5 –** Enrichment of SNP heritability, for additive effects and dominance effects (D), by bin for structural and evolutionary features in Ames/PHZ51+B47 and NAM/PHZ51, while accounting for enrichment by gene proximity by adding one bin in functional polygenic models for SNPs ≤ 1 kb from an annotated gene (Proximal). Gene+Rec.: recombination rate; Gene+MNase HS: chromatin openness; Gene+MAF: minor allele frequency; Gene+GERP: genomic evolutionary rate profiling score.

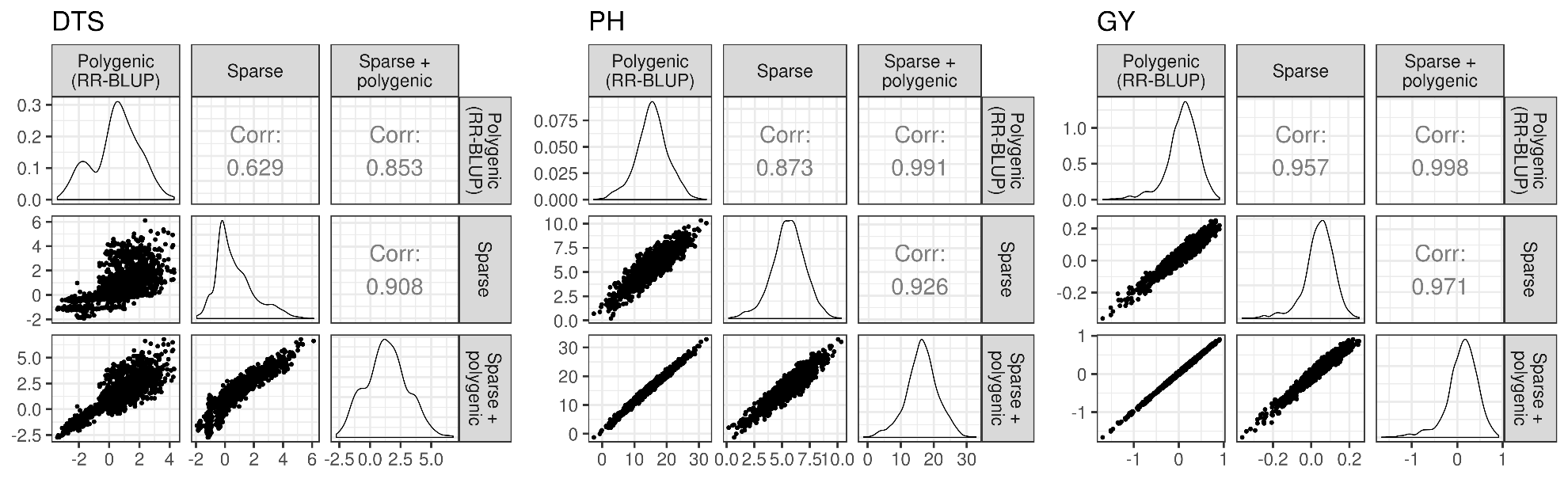

**Figure S6 ­–** Correlation and scatter plots for predicted genotype means in NAM/PHZ51 based on Ames/PHZ51+B47, for each trait (DTS, PH, and GY) and pair of effect types, in RR-BLUP (Polygenic) and BSLMM (Sparse and Sparse +polygenic).
